# *In vivo* RyR1 reduction in muscle triggers a core-like myopathy

**DOI:** 10.1101/2020.08.27.269647

**Authors:** Laurent Pelletier, Anne Petiot, Julie Brocard, Benoit Giannesini, Diane Giovannini, Colline Sanchez, Lauriane Travard, Mathilde Chivet, Mathilde Beaufils, Candice Kutchukian, David Bendahan, Daniel Metzger, Clara Franzini Armstrong, Norma B. Romero, John Rendu, Vincent Jacquemond, Julien Fauré, Isabelle Marty

## Abstract

Some mutations in the *RYR1* gene lead to congenital myopathies, through reduction in this calcium channel expression level, but the functional whole organism consequences of reduction in RyR1 amount have never been studied. We have developed and characterized a mouse model with inducible muscle specific *RYR1* deletion. Recombination in the *RYR1* gene resulted in a progressive reduction in the protein amount and was associated with a progressive muscle weakness and atrophy. Calcium fluxes in isolated muscle fibers were accordingly reduced. Alterations in the muscle structure were observed, with fibers atrophy, abnormal mitochondria distribution, membrane remodeling, associated with increase in the expression level of many proteins and inhibition of the autophagy process. This model demonstrates that RyR1 reduction is sufficient to recapitulate most features of Central Core Disease, and accordingly similar alterations were observed in muscle biopsies from Central Core Disease patients, pointing to common pathophysiological mechanisms related to RyR1 reduction.

## Introduction

One key step for skeletal muscle contraction is the intracellular calcium release performed by the sarcoplasmic reticulum calcium channel ryanodine receptor (RyR1) during the excitation-contraction coupling process. RyR1 is the core of the Calcium Release Complex (CRC), anchored both in the sarcoplasmic reticulum (SR) membrane and the T-tubule membrane, in a specific region of the muscle called the triad where these two membranes are in close apposition. Additional proteins are associated with RyR1 to form the CRC, including the voltage-activated plasma membrane calcium channel dihydropyridine receptor (DHPR) (Marty *et al.*, 1994), and regulatory proteins like the sarcoplasmic reticulum calcium binding protein calsequestrin (CSQ), or the sarcoplasmic reticulum membrane proteins triadin and junctin (Zhang *et al.*, 1997). Numerous mutations in the *RYR1* gene encoding the ryanodine receptor result in genetic diseases among which two core myopathies: Central Core Disease (CCD, OMIM#117000) and Multiminicore disease (MmD, OMIM#255320). The impact of *RYR1* mutations are either an alteration of the RyR1 calcium channel function (gain- or loss-of-function), a modification of the RyR1-DHPR coupling, or the reduction in the amount of RyR1 protein. In all cases, the consequence is a reduction in the amplitude of calcium release upon stimulation, resulting in muscle weakness (Dirksen and Avila, 2002; Bertzenhauser *et al.,* 2010; Lanner *et al.,* 2010; MacLennan and Zvaritch 2011; Marty and Fauré, 2016; Jungbluth *et al.,* 2018). To explore the various pathophysiology mechanisms associated with *RYR1* mutations, different mouse models have been created. They mostly reproduce amino acid substitutions in the RyR1 protein associated with mutations identified in patients in dominant (Y522S, I4898T; Chelu *et al.,* 2006; Zvaritch *et al.,* 2009) or recessive pathologies (T4706M+del; Q1970fsX16+A4329D) (Brennan *et al.,* 2019; Elbaz *et al.,* 2019*a*,*b*). These models have been useful to better understand some aspects of RyR1 physiology and to confirm the pathogenicity of patients’ mutations, but reveal mechanisms that are specific to individual or combination of RyR1 mutations.

In the present work we focused on the pathophysiology of the exclusive reduction in RyR1 protein expression in muscle, as this situation is frequently observed in patients affected with recessive congenital myopathy. To dissect the mechanisms related to a reduction in RyR1 amount we have developed and characterized a mouse model with a conditional and inducible *RYR1* knock out. In mice, the full deletion of the *RYR1* gene is lethal at birth (Takeshima *et al.,* 1994), and heterozygous deletion of a single *RYR1* allele results only in a modest 15% reduction in RyR1 protein, with no functional consequences (Cacheux *et al.,* 2015). In order to obtain a model with reduced amount of RyR1 specifically in skeletal muscles, we have developed a new mouse line RyR1^Flox/Flox^::HSA-Cre-ER^T2^ in which a decreased *RYR1* expression is induced by tamoxifen injection. We show that few weeks after induction, the amount of RyR1 protein reached 50% of initial level, and the mice developed a progressive myopathy, which recapitulates the main features observed in patients affected with Dusty Core Disease (a subgroup of Central Core Disease) and with a similar RyR1 reduction (Garibaldi *et al.,* 2019). This model with presents the same molecular and structural modifications as patients could therefore be a valuable tool for further pathophysiological mechanisms dissection and for therapeutic development.

## Results

### Generation of the RyR1^Flox/Flox^::HSA-Cre-ER^T2^ mouse model

A mouse line with loxP sites inserted on both side of exons 9-11 of the *RYR1* gene (so called RyR1^Flox/Flox^) was mated with the mouse line HSA-Cre-ER^T2^ expressing the tamoxifen dependent Cre-ER^T2^ recombinase (at heterozygous state) under the control of the human skeletal muscle α-actin gene (Schuler *et al.,* 2005), to create RyR1^Flox/Flox^::HSA-Cre-ER^T2^. Tamoxifen injection induces activation of the Cre-ER^T2^ recombinase selectively in skeletal muscle, which results in the deletion of exons 9-11 and disruption of the *RYR1* gene in skeletal muscle with a strict tamoxifen dependence (Figure 1A). At birth, RyR1^Flox/Flox^::HSA-Cre-ER^T2^ mice were normal, and at 2 months of age, once young adults, the recombination was induced by tamoxifen injection (from this time point, the recombined animals were called RyR1-Rec). Molecular and physiological consequences were further studied as a function of time, up to 105 days after induction of recombination (Figure 1B). Control animals (CTRL) are littermates RyR1^Flox/Flox^ (without the HSA-Cre-ER^T2^ transgene) injected with tamoxifen. Animals have been affected to the CTRL or the RyR1-Rec group depending on their genotype (with or without HSA-Cre-ER^T2^). The relative amount of RyR1 at the mRNA level was analyzed in quadriceps muscle every 15 days after recombination using RT-q-PCR in CTRL and in RyR1-Rec animals (Figure 1C). A rapid drop in RyR1 mRNA was observed in RyR1-Rec animals, with a low and stable level of 21% ± 4% of the initial value reached as soon as 3 days after tamoxifen injection. A reduction was observed in all of the tested muscles 75 days after recombination induction (*Quadriceps*, *tibialis anterior, EDL, soleus*, Supplementary Figure S1A). The amount of RyR1 protein was further analyzed, using quantitative Western blot in quadriceps muscle homogenates (Fig1D-E).

**Fig.1.**
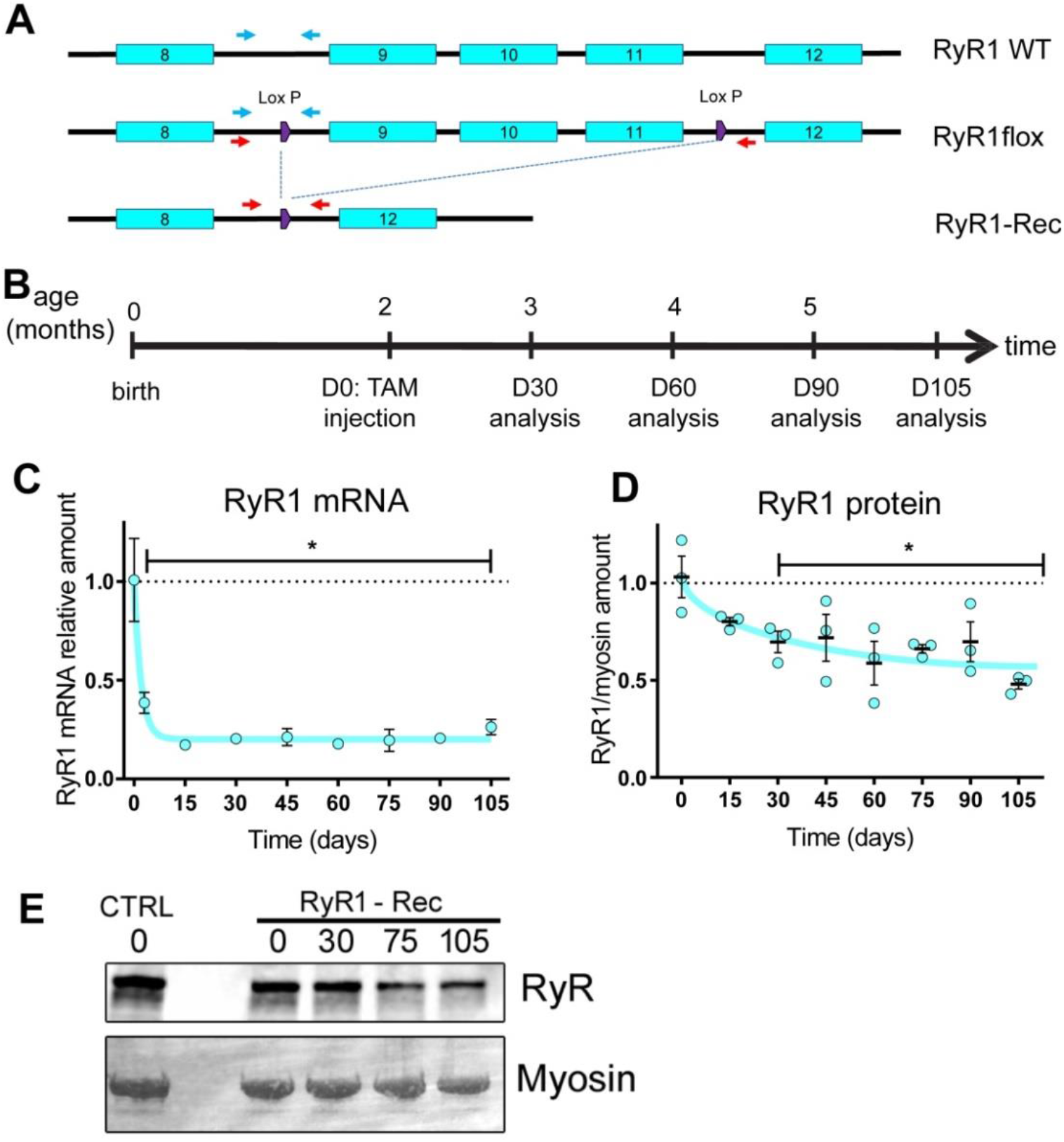
RyR1 mRNA and protein decrease after tamoxifen injection in the RyR1^Flox/Flox^::HSA-Cre-ER^T2^ mouse model. (A) LoxP sites were inserted on both sides of exons 9-11 in the RyR1 WT allele to create the RyR1-flox allele. After recombination, the RyR1-Rec allele is deleted with exons 9-11. The localization of the primers (blue and red) to allow discrimination between the different alleles is presented. (B) The animals were injected with tamoxifen to induce the recombination at 2 months of age (D0), and were analyzed at variable times thereafter. (C) The relative amount of mRNA compared to beta-actin, HPRT and GAPDH as reference genes were evaluated using RT-q-PCR in quadriceps muscles of n=3-6 different mice at each time point, and is presented as mean ± SEM for each time. The amount in CTRL littermate was set to 1. The quantification was performed using the ΔΔCt method. Statistical analysis: One way ANOVA with Holm-Sidack’s test for multiple comparisons (D) The relative amount of RyR1 compared to myosin heavy chain was evaluated using quantitative Western blot in quadriceps homogenates of n=3 different mice at each time point. Individual data and mean ± SEM are presented. The initial amount was set to 1. (E) Representative Western blot of RyR1 on CTRL and RyR1-Rec quadriceps homogenates at different time points, using myosin heavy chain as a control of protein amount. Statistical analysis: One way ANOVA with Holm-Sidack’s test for multiple comparisons.

This amount was progressively reduced and reached about 50% of initial value after 105 days. No further reduction in RyR1 protein was observed on longer times (data not shown). The amount of RyR1 protein was quantified in different muscle homogenates 75 days after recombination. A reduction was observed in all the assayed muscles, although the remaining amount slightly differed between the muscles, the highest being in *interosseous* (64% ± 5%) and the lowest in *soleus* (33% ± 5%) (Supplementary Figure S1B).

### RyR1-Rec mice show progressive reduction in muscle and body weights and in muscle strength

The consequences of RyR1 reduction were first studied at the whole animal level. Initially at the same weight, the CTRL animals slowly gained weight from 24.4 ± 0.4g to 30.6 ± 0.8g at D90 whereas the RyR1-Rec animals progressively lost weight to 20.6 ± 1.3g at D90 (Figure 2A). This is the result of a weight loss observed in all the muscles (Supplementary Figure S1C). To test the overall muscle performance after RyR1 reduction, the animals were subjected to two different strength tests. They were first allowed to hang gripping a cross-wired surface with all four paws up to 5 min, the latency to fall reflecting the muscle force. The grip test performed every week during 105 days (Figure 2B) showed that RyR1-Rec animals started losing strength 20-30 days after tamoxifen injection, when the amount of RyR1 was about 75-80% of its initial value, and were unable to hang on the grid about 75 days after tamoxifen injection, as RyR1 amount had reached the level of about 60%. Muscle strength was also assessed by a 6 min noninvasive electrostimulation protocol of the *gastrocnemius* muscle coupled to anatomic magnetic resonance imaging (MRI) under general anesthesia. During the electrostimulation protocol of CTRL animals 30 days after tamoxifen injection (Figure 2C, D30) a typical exercise profile was recorded, with a stable initial muscle strength which slowly declined as muscle fatigued.

**Fig. 2.**
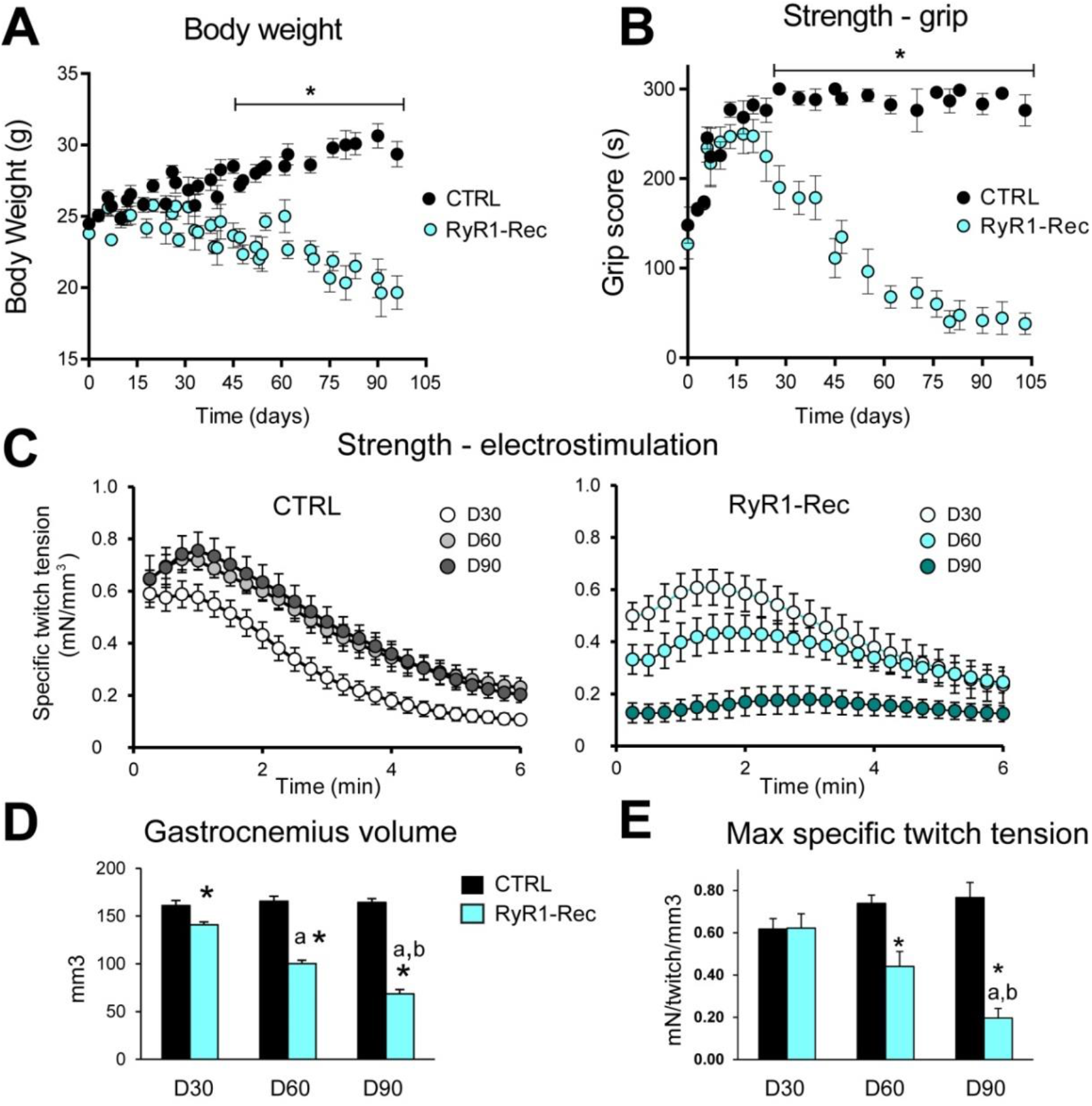
RyR1-Rec mice show progressive reduction in muscle and body weights and in muscle strength. (A) Body weight and (B) Muscle strength estimated as a function of time after recombination using a grip test in which the time the animals can hang upside down on a grid up to 5 min (300s). Data are means ± SEM of n= 10-15 animals in each group, *\* p<0.05* Student’s t-test with Holm-Sidack’s correction for multiple comparisons (C) Recording of the tension developed by gastrocnemius during a 6 min electrostimulation protocol at 2Hz. Longitudinal study of the same animals (CRTL, left panel, and RyR1-Rec, right panel) at different times after recombination, 30 days (D30), 60 days (D60) and 90 days (D90). Data are means ± SEM of n=8 CRTL and n=6 RyR1-Rec animals. (D) *Gastrocnemius* muscle volume for CTRL (n=8) and RyR1-Rec (n=6) mice at 30, 60 and 90 days after recombination. Data are means ± SEM. Statistical analysis: post-hoc LSD Fisher test following two-way repeated measures ANOVA, * significantly different from CTRL at the same time, ^a^ significantly different from D30 in the same group, ^b^ significantly different from D60 in the same group. (E) Maximal specific twitch tension (maximal twitch tension normalized to the gastrocnemius volume). Statistical analysis: post-hoc LSD Fisher test following two-way repeated measures ANOVA * significantly different from CTRL at the same time, ^a^ significantly different from D30 in the same group, ^b^ significantly different from D60 in the same group.

In the CTRL group at D60 and D90 the exercise profile showed improved muscle force when the animals got older (Figure 2C). In contrast, the performance of RyR1-Rec animals, similar to CTRL at D30, deteriorated with age. RyR1-Rec animals were unable to perform the exercise anymore at D90 (Figure 2C, RyR1-Rec D90). *Gastrocnemius* volume measured using MR images (Figure 2D) remained stable in CTRL animals from D30 (161±5 mm^3^) to D90 (164±4 mm^3^). In contrast, in RyR1-Rec mice, *gastrocnemius* volume was significantly smaller than in CTRL mice since D30 (141±3 mm^3^, *p=0.003*), and its volume was further reduced down to 100±3 mm^3^ at D60 (*p<0.001* compared to CTRL at the same age) and 69 ± 4 mm^3^ at D90 (*p<0.001* compared to CTRL at the same age). The maximal specific twitch tension (absolute twitch tension normalized to muscle volume) confirms that both groups presented similar muscle strength at D30 (Figure 2E), but RyR1-Rec animals’ strength dramatically declined whereas it increased in CTRL animals as they got older. Additionally, the changes in *gastrocnemius* muscle bioenergetics during the electrostimulation were assessed noninvasively at D60 using *in vivo* 31-phosphorus (^31^P) MR spectroscopy. Whereas the basal intramyofibrillar pH did not differ between both groups (Supplementary Figure S2A), the extent of acidosis at the end of the exercise was lower (*p=0.016*) in RyR1 Rec animals (Supplementary Figure S2B), thereby suggesting that glycolytic flux in exercising muscle was reduced in these animals. Moreover, the time constant of post-exercise phosphocreatine resynthesis (τPCr) was significantly shorter (*p=0.047)* in RyR1-Rec mice (Supplementary Figure S2C-D), reflecting an improved *in vivo* mitochondrial function. Indeed, PCr synthesis during the post-exercise recovery period relies exclusively on oxidative ATP synthesis, thus τPCr is considered as an index of the oxidative phosphorylation capacity (Kemp *et al.,* 1995). Overall, these results indicate that RyR1 progressive reduction is associated with a progressive loss of weight and a drop in muscle strength.

### The physiological properties of isolated fibers are compromised in RyR1-Rec mice

The alteration of muscle function was further characterized at the single muscle fiber level using a combination of whole-cell voltage-clamp and confocal imaging (Lefebvre *et al.,* 2014; Kutchukian *et al.,* 2016). The T-tubule network was imaged by staining the plasma membrane with di-8-anepps. No qualitative difference in the network (Figure 3A, left) or quantitative difference in the mean T-tubule density between CTRL and RyR1-Rec fibers were observed, beside a slight but significant shortening of the mean resting sarcomere length (Figure 3A, graphs on the right). The voltage-activated Ca^2+^ influx through the DHPR (Figure 3B-D) and of the Ca^2+^ release flux through RyR1 (Figure 3E-I) were simultaneously assessed. As compared to CTRL fibers, RyR1-Rec fibers exhibited voltage-activated DHPR Ca^2+^ currents of similar time course (Figure 3B) but reduced density, as shown by the peak current density *vs* voltage in the two groups (Figure 3C), which translated into a 30 % reduction in maximum conductance (G_max_) with no associated change in the other parameters of the current *vs.* voltage relationship (Figure 3D). The voltage dependence of SR Ca^2+^ release was assessed from line-scan imaging of the Ca^2+^-sensitive dye rhod-2. Line-scan images from the RyR1-Rec fibers were qualitatively similar to those from CTRL fibers (Figure 3E), showing a rapid rise in fluorescence upon T-tubule membrane depolarization, spatially homogeneous along the scanned line. The rate of SR Ca^2+^ release (Figure 3G), calculated from the changes in rhod-2 fluorescence elicited by depolarizing pulses of increasing amplitude (Figure 3F), exhibited a similar time course in RyR1-Rec fiber as in CTRL fibers, but importantly, peak values were reduced in RyR1-Rec. Fitting of the peak rate of SR Ca^2+^ release *vs* voltage (Figure 3H-top graph) indicated that the maximum rate of Ca^2+^ release was reduced by 25% in the RyR1-Rec group (Figure 3I, Max), the slope factor (*k*) was also slightly but significantly reduced, whereas the voltage of mid-activation (*V_0.5_*) was unchanged.

**Fig.3.**
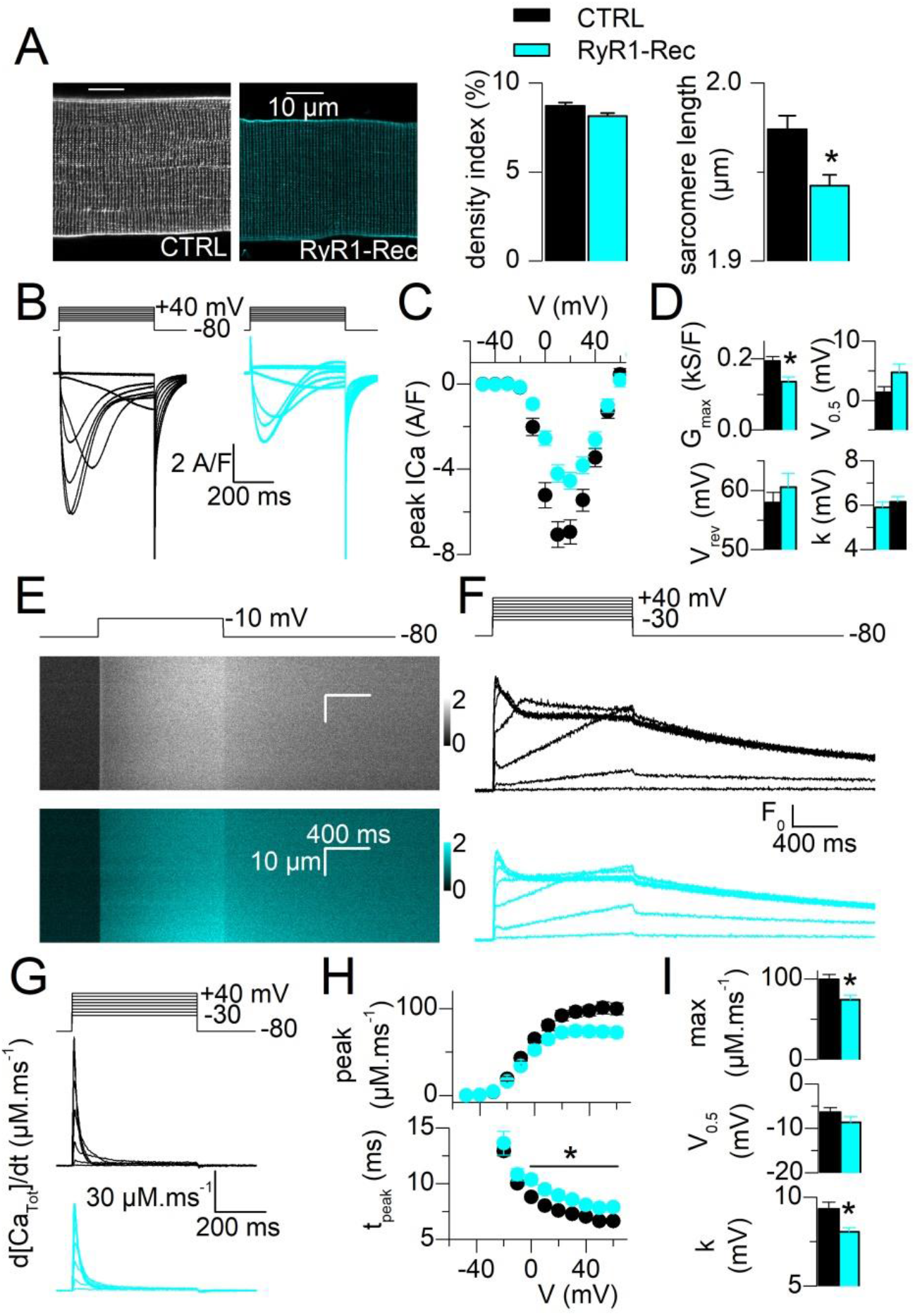
Excitation-contraction coupling in single isolated muscle fibers is altered. All values are means ± SEM. Statistical significance was determined using a Student’s t-test. (A) Representative confocal images of T-tubule network stained with di-8-anepps from a CTRL and a RyR1-Rec fiber, allowing evaluation of T-tubule density and sarcomere length, performed in 42 CTRL fibers and 41 RyR1-Rec fibers (4 mice in each group). (B) Representative DHPR Ca^2+^ current from a CTRL and a RyR1-Rec fiber in response to 0.5s-long depolarizing steps to the indicated levels (10 mV increment). (C) Voltage-dependence of the peak DHPR Ca^2+^ current density. (D) Parameters obtained from the fit of curves *(C)* in 29 CTRL fibers and 30 RyR1-Rec fibers (5 mice in each group), (cf Supp. Methods). (E) Representative *x,t* images of rhod-2 fluorescence (a.u.) from a CTRL and a RyR1-Rec fiber stimulated by a voltage-clamp depolarizing pulse to −10 mV. (F) Representative line-averaged rhod-2 Ca^2+^ transients from a CTRL and from a RyR1-Rec fiber in response to voltage-clamp pulses to the indicated levels. (G) Rate of SR Ca^2+^ release calculated from the curves shown in *(F).* (H) Voltage-dependence of the peak rate of SR Ca^2+^ release (top) and of the time-to-peak rate of SR Ca^2+^ release (bottom). (I) Parameters obtained from fitting the peak rate of SR Ca^2+^ release *vs* voltage relationship with a Boltzmann function in each fiber (cf Supplementary Method). Data are from the same fibers as in *(B-D)*.

A slight (1.3ms in average) but significant increase in the time-to-peak rate of SR Ca^2+^ release in the RyR1-Rec fibers (Figure 3H, bottom graph) was observed. No modifications were observed in the cytosolic Ca^2+^ removal capabilities of the fibers (SERCA pump function), measured with the low-affinity Ca^2+^-sensitive dye fluo-4 FF in non-EGTA-buffering conditions (Supplementary Figure S3). Overall, these results demonstrate that the 35% reduction in RyR1 amount in *interosseous* is associated with a 30% reduction in calcium current through DHPR and a 25% reduction in calcium flux through RyR1.

### RyR1 reduction affects the skeletal muscle structure

To better understand the basis of these functional alterations associated with a decrease in RyR1, muscle structure was studied on RyR1-Rec animals at D75 after recombination, and compared to CTRL animals. *Extensor digitorum longus* (EDL), a fast twitch muscle, *soleus*, a slow twitch muscle and *tibialis anterior* (TA), a mixed muscle were analyzed using hematoxylin-eosin, NADH and Gomori trichrome stainings. Stainings of TA, presented in Figure 4A, show abnormal muscle structure in RyR1-Rec animals, without fibrosis or central nuclei, but with fiber atrophy, affecting both type I and type II fibers (Table 1).

**Table 1:** Analysis of muscle fibers’ diameter

| | CTRL<br>diameter<br>($\mu\text{m}$) | SEM | Nb<br>fiber | RyR1-Rec<br>diameter ($\mu\text{m}$) | SE<br>M | Nb<br>fiber | Student<br>t-test | %CTR<br>L |
| --- | --- | --- | --- | --- | --- | --- | --- | --- |
| EDL type I | 23,6 | 0,9 | 700 | 17,2 | 0,7 | 700 | $p<0.001$<br>* | 73 |
| EDL type II | 40,0 | 1,6 | 600 | 32,5 | 1,2 | 700 | $p<0.001$<br>* | 81 |
| TA type I | 33,7 | 1,9 | 300 | 20,0 | 1,2 | 300 | $p<0.001$<br>* | 59 |
| TA type II | 46,8 | 2,7 | 300 | 35,6 | 2,1 | 300 | $p<0.001$<br>* | 76 |

The diameter of slow (type I) and fast (type II) fibers was quantified in *Extensor digitorum longus* (EDL) and *tibialis anterior* (TA) sections of CTRL and RyR1-Rec animals at D75 after NADH-TR staining. Hundred fibers were measured for each muscle. Between 3 and 7 mice in each group. \* represent significant data.

**Fig.4.**
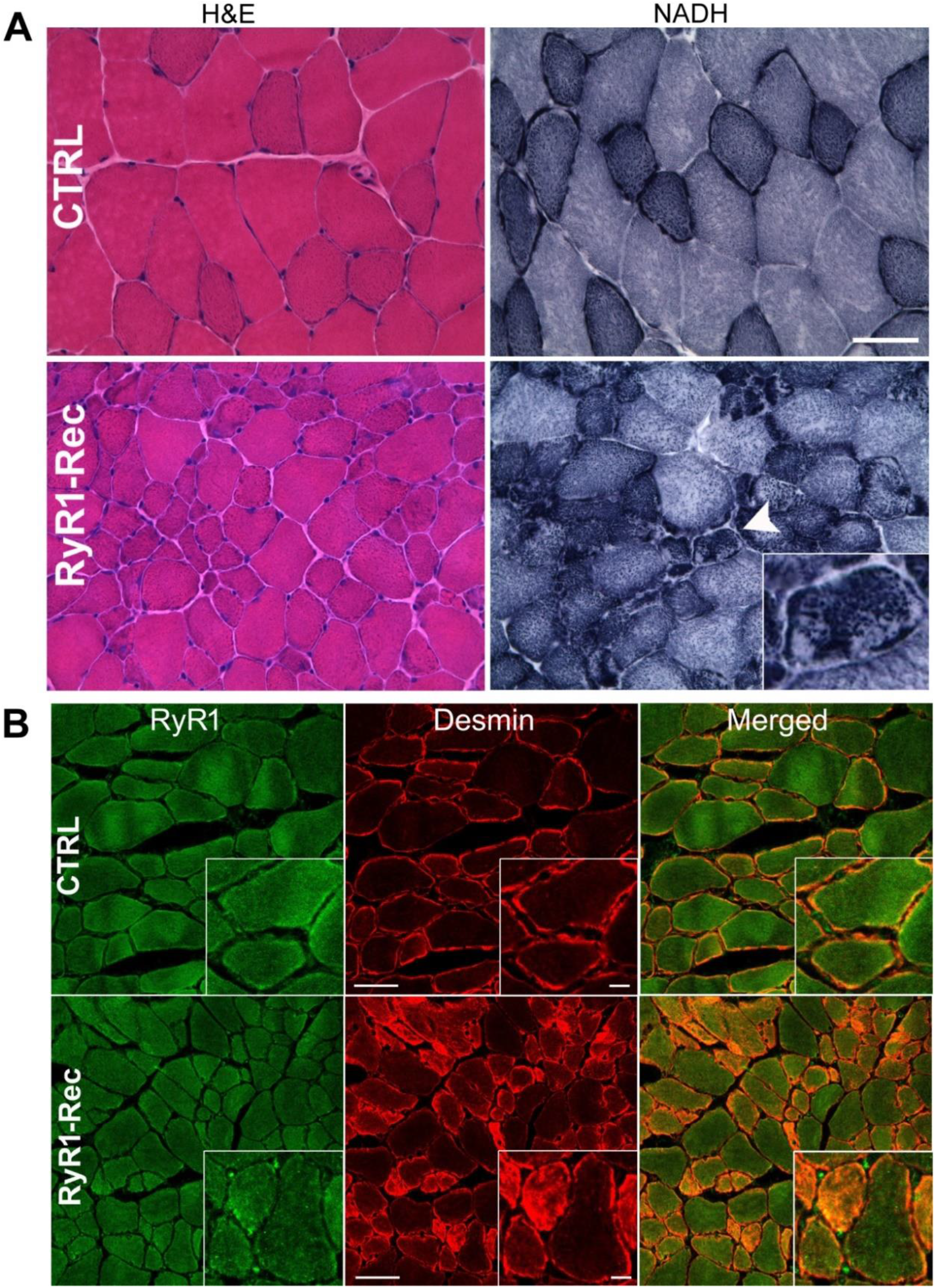
Histological and immunofluorescent analyses show major defects in muscles. (A) *TA* Transversal section from CTRL and RyR1-Rec mice at D75 were stained with hematoxylin/eosin (H&E) and NADH. The mitochondria localization (NADH staining) is inhomogeneous in RyR1-Rec fibers, specifically in the small dark type I (slow) fibers with high mitochondria content (arrow head and inset). The nuclei (blue dots with H&E staining) are at the periphery of the fibers, and no evidence of regenerative fiber can be seen. Bar 50μm. (B) Transversal TA sections from D75 CTRL and RyR1-Rec mice were stained with antibodies against RyR1 (green) and desmin (red). Bar 50μm, and 10μm in inset.

A mitochondria disorganization characterized by local accumulation/depletion in adjacent regions of the same fiber (arrowhead and inset Figure 4A) and red staining accumulation observed with Gomori trichrome (Supplementary Figure S4) was also observed. Fiber atrophy was observed in both fiber types in EDL, although the reduction was slightly more important in type I fibers (Table 1). Transversal TA section of CTRL and RyR1-Rec mice were stained with antibodies against RyR1 and desmin. No modification was observed in the RyR1 distribution between type I and type II fibers in RyR1-Rec TA sections, indicating a similar RyR1 reduction in all fiber types (Fig. 4B). Surprisingly, an important modification was observed in desmin distribution, with a huge increase in the small type I fibers (Fig. 4B): in CTRL TA sections, all fibers presented a similar peripheral staining, whereas the small type I fibers in RyR1-Rec TA sections were heavily stained both at the fiber periphery and in the cytosol (inset Fig. 4B).

Triads organization was further studied using immunofluorescent labelling of isolated EDL fibers (Figure 5A). The labeling of alpha-actinin (as a marker of the Z-line), of triadin and of RyR1 (as triad markers) on RyR1-Rec EDL fiber demonstrated that the reduction in RyR1 amount resulted in a generalized disorganization of the fiber with abnormal triads localization and disruption of the Z-line, but triadin and RyR1 were still partially colocalized. Although triads remained on both sides of the Z-line, the Z-line did not form a straight line as in CTRL and instead presented many micro-disruptions (inset Figure 5A). Muscle ultrastructure was analyzed using electron microscopy on RyR1-Rec EDL fibers at D75 (Figure 5B). Some regions had a relatively preserved structure (Figure 5B, left fiber), whereas some adjacent regions were highly disorganized, with disruption of sarcomere regular organization, mitochondria disorganization, and the presence of numerous stacks of membrane (Figure 5B, arrows, enlarged in inset), previously called multiple triads (Paolini *et al.,* 2007, Garibaldi *et al.,* 2019).

**Fig. 5.**
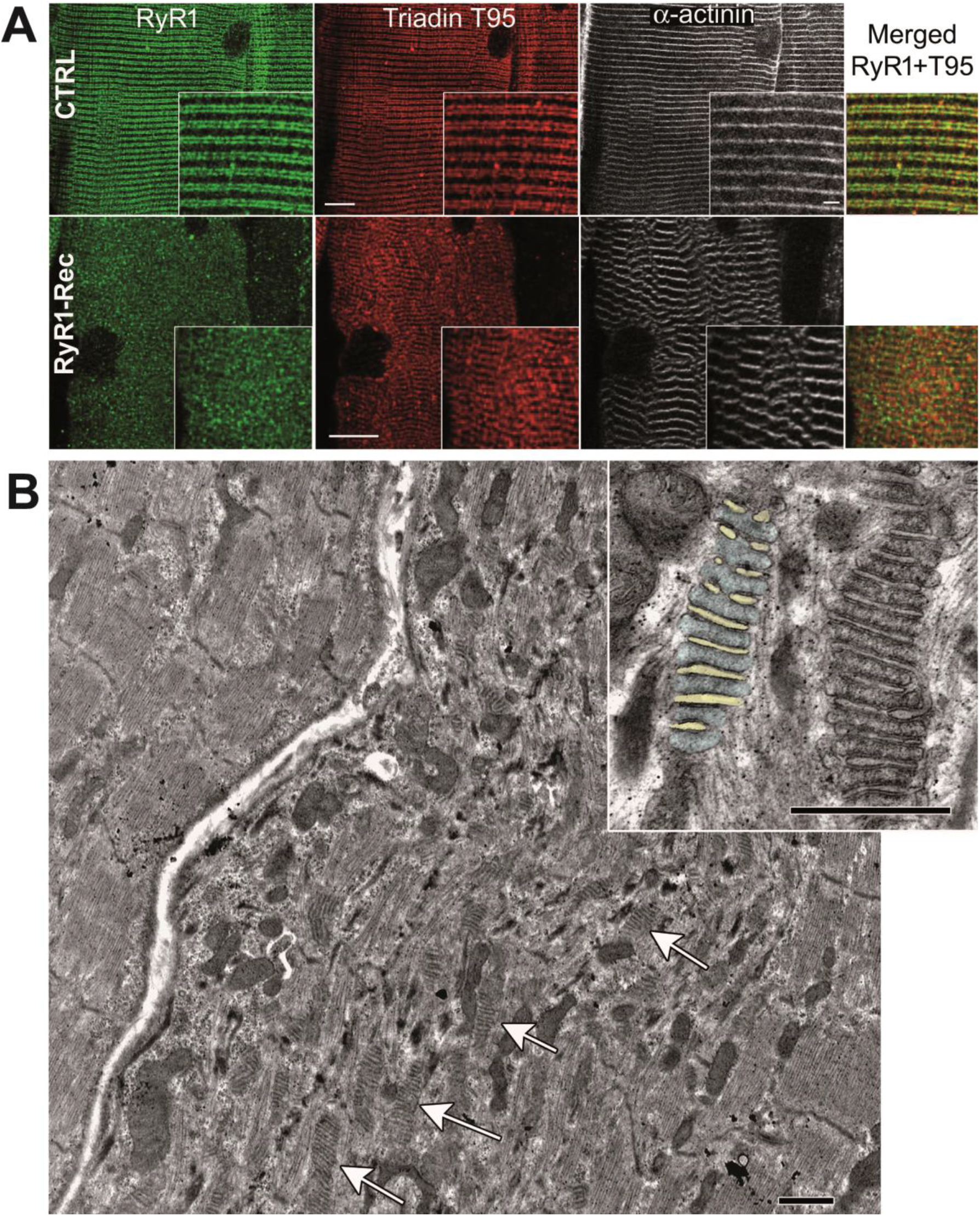
A generalized structural disorganization is observed in EDL muscle fibers. (A) EDL fibers from D75 CTRL and RyR1-Rec mice were stained with antibodies against RyR1 (green), triadin (red) and alpha-actinin (white). Bar 10μm and 2μm in inset. (B) Electron microscopy analysis of longitudinal *EDL* section of D75 RyR1-Rec mice. The upper left fiber has a normal structure, the adjacent lower fiber is extremely disorganized, with large stacks of membranes (up to 15 stacks, arrows) extending on few μm long. The inset presents two multiple triads, on the left one the membranes corresponding to T-tubules have been colored with light yellow and the SR sheets with light blue to allow a better visualization. Some electron dense material can be seen within the SR segment, all along the contact site with the adjacent T-tubule, which could correspond to accumulation of proteins. Bars 1μm.

The morphological features of those multiple triads compared to the normal single triads are presented in Table 2, and additional examples are presented in supplementary Figure S5. Precise quantification of putative T-tubule width, putative SR width and intermembrane (T-tubule/SR) space (supplementary Figure S5) demonstrated a huge increase in the T-tubule mean size (two-fold increase, reaching 202% of the T-tubule size in normal triad) in those multiples triads, whereas the SR mean width was only slightly increased (+10%) and the intermembrane space was not modified. Those results point to a generalized structural disorganization of RyR1-Rec mice muscles associated with membranes remodeling.

**Table 2:** Morphological characterization of single and multiple triads

|  | Single (CTRL) |  |  | Multiple (RyR1-Rec) |  |  | Student t-test | % CTRL |
| --- | --- | --- | --- | --- | --- | --- | --- | --- |
|  | Mean | SEM | n | Mean | SEM | n |  |  |
| TT width (nm) | 26,0 | 0,5 | 140 | 52,7 | 1,6 | 135 | $p<0.001$<br>* | 202 % |
| SR width (nm) | 77,5 | 0,8 | 239 | 85,2 | 1,2 | 107 | $p<0.001$* | 110 % |
| SR-TT space (nm) | 20,6 | 0,3 | 135 | 20,5 | 0,4 | 138 | 0,9 | 100 % |

The size of T-tubules, SR and intermembranes space thickness were measured as presented on supplementary figure S5. \* represent significant data.

### RyR1 reduction is associated with modification in the amount of numerous proteins

The consequences of RyR1 reduction were studied at the molecular level using quantitative Western blot on quadriceps muscle of CTRL or RyR1-Rec mice (8 animals in each group) at D75 after tamoxifen injection (Figure 6). The relative amount of numerous proteins involved directly or indirectly in calcium handling was estimated, using the total amount of proteins determined by stain free evaluation as reference (Gürtler *et al.,* 2013). No difference in the quantification of RyR1 was observed between this method and the use of myosin heavy chain as a reference protein (comparison between Figure 1 and Figure 6). No significant modification between CRTL and RyR1-Rec muscles was observed in the amount of the triadin isoform T95, the calcium binding protein CSQ1, the Ca^2+^-ATPase SERCA, the mitochondrial FoF_1_-ATPase, and the alpha 1 subunit of DHPR (Fig. 6A and B).

**Fig.6.**
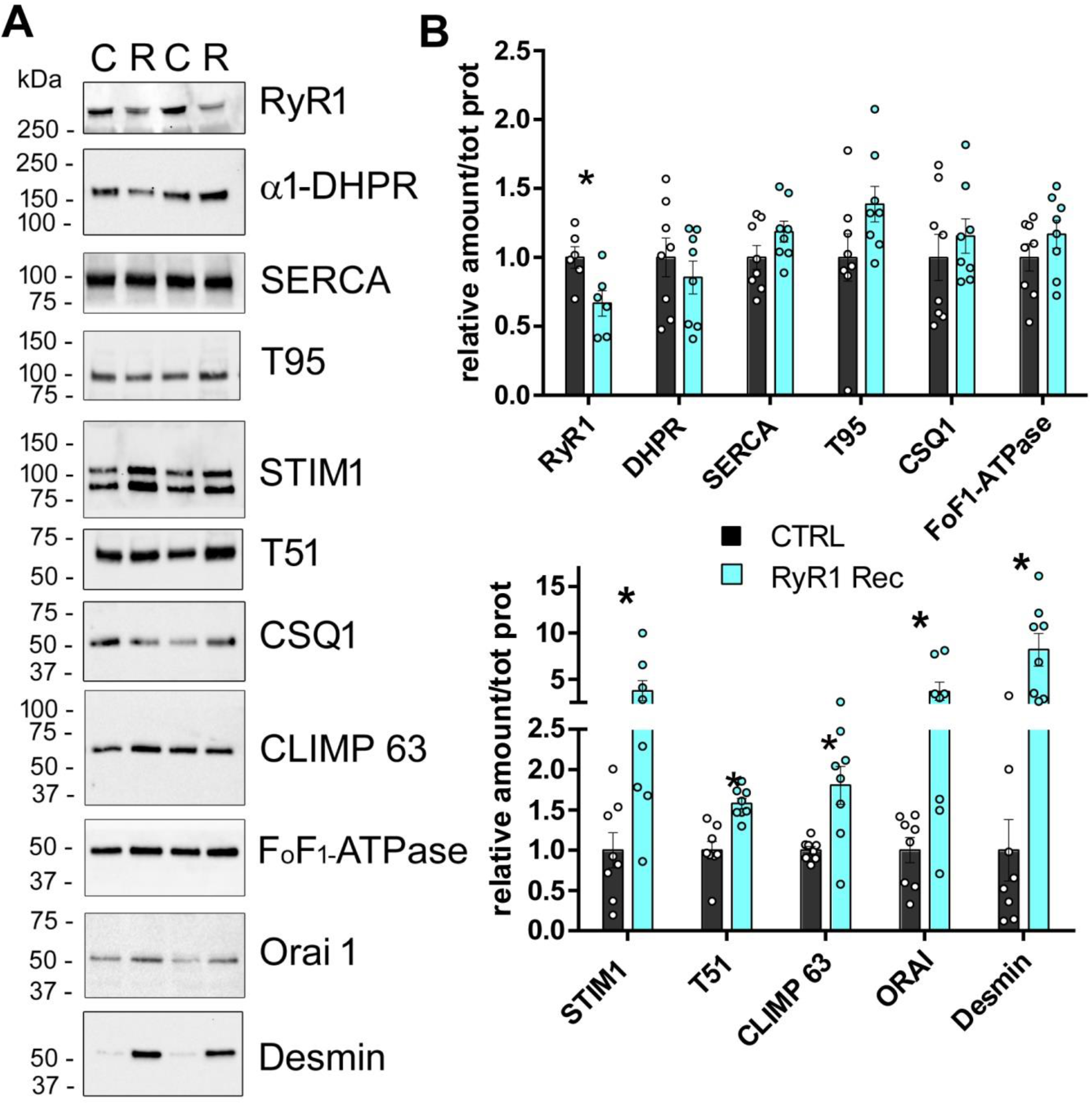
Quantitative Western blot analysis of the proteins expressed in quadriceps muscles of D75 mice demonstrate an increase in expression of many proteins. (A) Representative Western blots for each protein on D75 quadriceps homogenates from 2 different CTRL (C) and RyR1-Rec (R) mice. (B) Quantification of the amount of each protein normalized to the total amount of proteins on 8 different animals in each group (CTRL, back bars and RyR1-Rec, blue bars). The value for each animal is the mean of at least 3 blots. All the data are presented as mean ± SEM. The mean value in the CTRL group was set to 1 for each protein. Statistical analysis: t-test with Holm-Sidak method for multiple comparisons.

In contrast, an increase was observed in the amount of STIM1 (x3.7±1, *p=0.02*), the triadin isoform T51 (x1.6±0.1, *p<0.001*), CLIMP63 (x1.8±0.2, *p=0.004*), and ORAI1 (x3.7±1, *p=0.017*). As modification in the localization of the structural protein desmin was observed using immunofluorescent labeling (Fig. 5B), desmin expression level was also quantified and a huge increase in desmin amount was observed (x8.2±1.2, *p=0.001*). The localization of CLIMP63 and STIM1 in RyR1-Rec EDL fibers was analyzed using immunofluorescent labeling, and no major modification was observed (Supplementary Figure S6). Therefore RyR1 protein reduction was associated with an increase in different proteins involved in calcium regulation or in muscle architecture.

### Autophagy is altered as a result of RyR1 reduction

Alteration in autophagy has been observed in some myopathies (Durieux *et al.,* 2012; Al Qusairi *et al.,* 2013). In order to identify the mechanisms leading to the muscle atrophy subsequent to RyR1 reduction, we looked for modifications in the autophagy flux in RyR1-Rec muscles. The amount of LC3 II, a membrane protein of the autophagosomes, was evaluated with quantitative Western blot (Figure 7A and B), and an increase in LC3 II was observed (172% ± 32%, *p*=0.03). This increase in autophagosomes could reflect either an increase in formation of autophagosomes (due to an increase in autophagy process) or a decrease in their fusion with lysosomes (and therefore an inhibition of autophagy degradation). The precise nature of autophagy flux modification was further analyzed by quantification of p62, a well-known protein specifically degraded via autophagy, which amount was also significantly increased (Fig. 7A-B, 172% ± 22%, *p=0.006)*). The associated increase in LC3 II and in p62 in RyR1-Rec muscle compared to control suggested that RyR1 reduction was therefore associated with inhibition of the autophagy flux. Moreover, an increase in the activation (phosphorylation) of two inhibitors of autophagy, mTOR and S6 protein (Fig. 7B-C), was observed (P-mTOR/mTOR increase by 174% ± 17%, *p=0.01;* P-S6/S6 increase by 320 % ± 50%, *p<0.001)*. The increase in the phosphorylation of these two autophagy inhibitors in RyR1-Rec animals compared to controls further pointed towards an inhibition of autophagy responsible for the muscle atrophy in RyR1-Rec animals.

**Fig.7.**
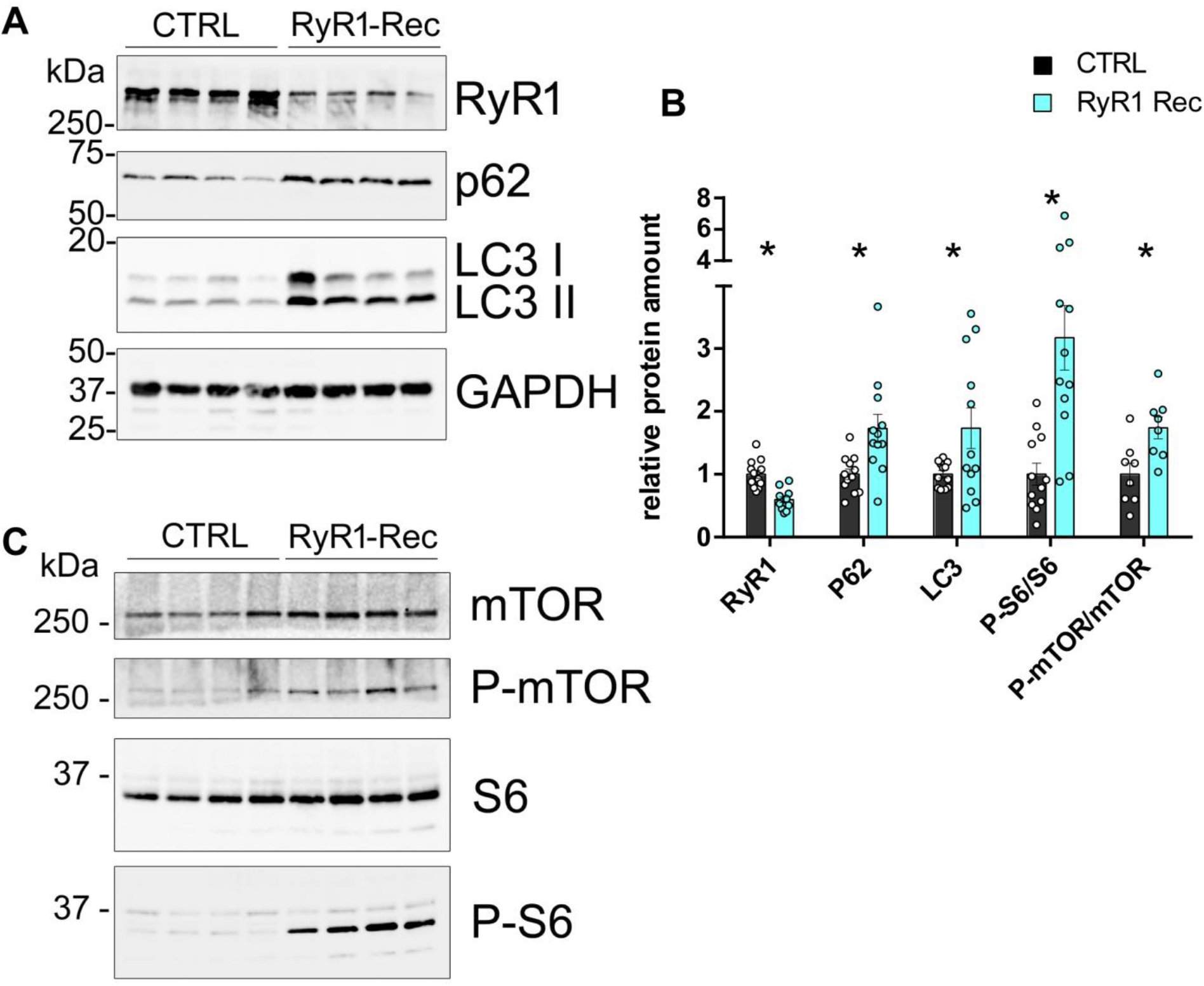
Autophagy is inhibited in RyR1-Rec D75 mice quadriceps muscle. (A) and (C) Representative Western blot on 4 CTRL and 4 RyR-Rec quadriceps muscle homogenates. (B) Quantification of the amount of protein normalized to the amount of GAPDH in 8 to 12 mice in each group (CTRL, black bars; RyR1-Rec, blue bars). For S6 and mTOR, the values are presented as ratio of the phosphorylated/non-phosphorylated protein. The value for each animal is the mean of at least 3 blots. All the data are presented as mean ± SEM. The mean value in the CTRL group was set to 1 for each protein. Statistical analysis: t-test with Holm-Sidak method for multiple comparisons.

### Human patients’ biopsies exhibit similar defects to those observed in RyR1-Rec mice

In a recent study on a large cohort of patients with a recessive congenital myopathy (Garibaldi *et al.,* 2019), we have identified the presence of atypical structures, called “dusty cores”, in more than half of the patients with a reduction in RyR1 amount. They were observed by multiple staining and differ from the classical central core (well defined regions devoid of oxidative stain) by their poorly defined borders, their focal myofibrillar disorganization, a reddish-purple granular material deposition at Gomori trichrome staining and blended area with decreased or/and increased enzymatic activity at oxidative stains (Garibaldi *et al.,* 2019). These structural alterations mirror the modifications observed in our mouse model (Figure 4). We therefore further compared our new mouse model and muscle biopsies from patients with a RyR1 protein reduction and “dusty cores”. Using quantitative Western blot, the amount of the proteins which were clearly modified in our mouse model was also estimated in 5 patients with a recessive Dusty Core disease (two RyR1 mutations resulting in RyR1 protein reduction - Figure 8A). Control biopsies of different age (between 3.5 and 64 years) were used as reference. A large reduction in RyR1 protein was observed in all the patients (mean expression level 19.8 ± 2.2% of CTRL, *p<0.001*). Moreover, an increase in CLIMP63 and desmin was also observed. The reduction in RyR1 was associated with an important increase in CLIMP 63 (mean expression level increase 5 folds, 516 % ± 220%). In addition, the expression level of desmin was also drastically increased in patients compared to control (mean expression level increase 240 folds, 24 170 % ± 7 635 %). Using electron microscopy to analyze patient biopsy’s ultrastructure, many stacks of membranes were observed in the disorganized regions (Figure 8C, arrows) of all the patients. These regions, similar to the stacks observed in RyR1-Rec muscle (Figure 4B), pointed to a common mechanism, both in mice and in human, leading to the formation of those structures as a result of RyR1 reduction. With identical modifications as the ones in Dusty Core Disease patients, this model constitutes thus a relevant model to study the pathophysiological mechanisms.

**Fig. 8.**
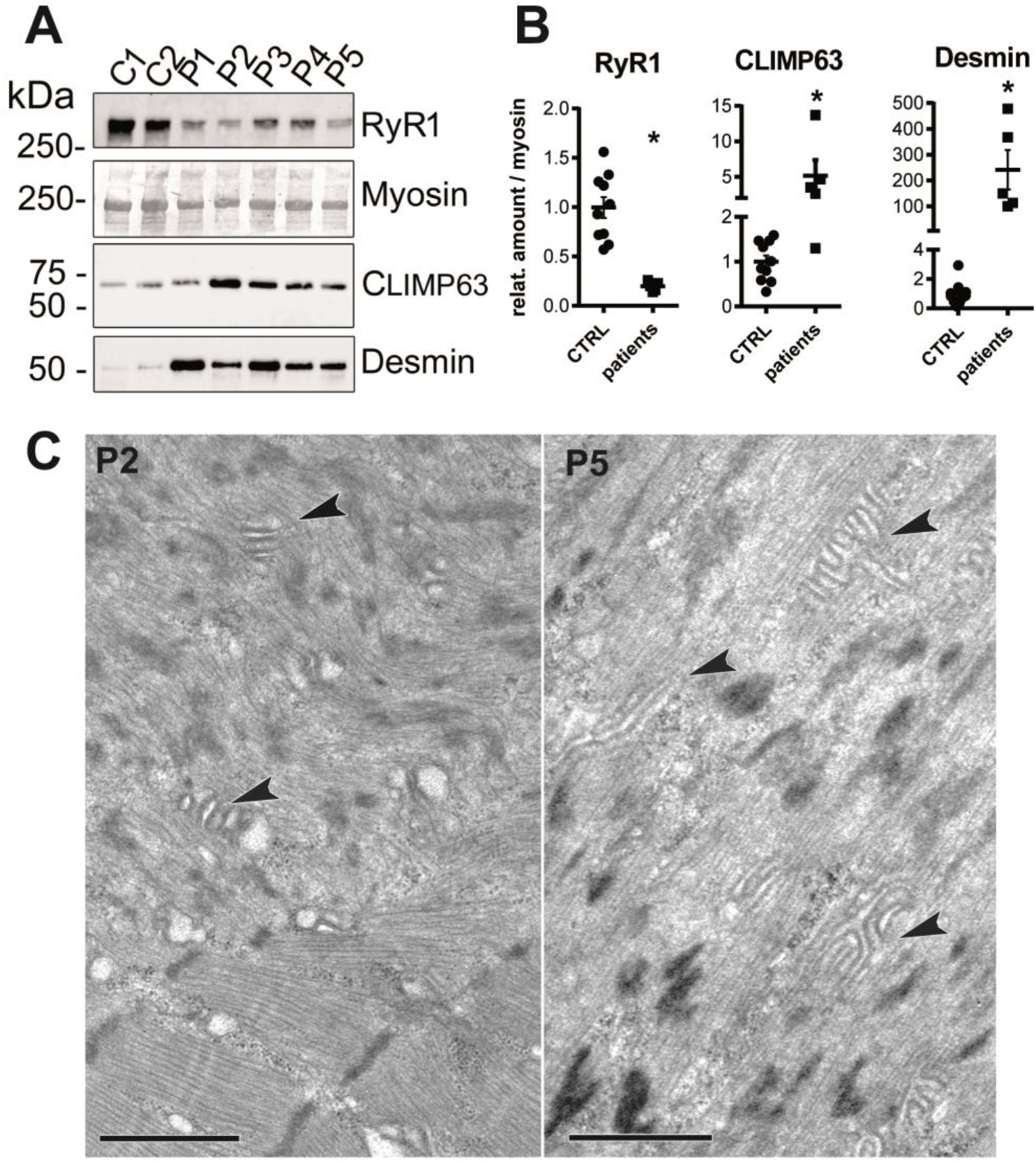
Analysis of human patients’ biopsies reveals similar defects to those observed in RyR1-Rec mice. (A) Representative Western blot performed on muscle homogenate from human biopsies. C1: control, 23 years, female; C2: control, 3.5 years, male; P1: CCD (Dusty core), mutations p.M2423K+p.R2441*, 43 years, male; P2: CCD (Dusty core), mutations p.T4709M + p.R1409*, 4 years, male; P3: CCD (Dusty core), mutations p.[Ile1571Val; Arg3366His; Tyr3933Cys] + p.Val788Cysfs*96, 25 years, female; P4: CCD (Dusty Core), mutations p.R2140W+p.L4828R, 9 years, female; P5: mutations p.M4000del+ p.Met2312Cysfs*118, 28 years, female. (B) Quantification of protein amount, normalized to myosin (for RyR1, CLIMP63 and Desmin) or to GAPDH (for p62). Data are presented as mean ± SEM of 10 controls samples (CTRL) and of 5 patients’ samples (patients). The value for each patient is the mean of at least 2 Western blots. The mean value for each protein in CTRL samples was set to 1. RyR1 expression level compared to controls: P1- 26 ± 6%, P2- 14 ± 6%, P3- 20 ± 3%, P4- 23 ± 3%, P5- 16 ± 2%. CLIMP63 expression level compared to control: P1- 130 % ± 12%, P2- 1 372 % ± 385%, P3- 466 % ± 92%, P4- 262 % ± 35%, P5- 350 % ± 85%). Desmin expression level compared to control: P1- 47 789 % ± 6097 %; P2- 10 052 % ± 2 957 %; P3- 36 745 % ± 7 150 %; P4- 14 951 % ± 5 584 %; P5- 11 307 % ± 2 858 %. Student t-test RyR1 *p<0.001*, CLIMP63 *p= 0.016*, Desmin *p<0.001* (C) Electron microscopy pictures obtained during the course of the diagnosis, presenting multiples stacks of membranes in the disorganized core region of the biopsies of patients P2 and P5. Similar structures were identified in the muscle biopsies of the five patients. Bar 1μm.

## Discussion

We present here the first model of mice with a clear myopathy phenotype due to the exclusive reduction in the expression of the *RYR1* gene. The recombination of the *RYR1* gene in this inducible and muscle specific RyR1-KO mouse line was performed at the adult stage, and induced a progressive reduction in RyR1 protein amount in adult and well-structured muscles, thus avoiding possible developmental effects of *RYR1* knockdown. Although the production of *RYR1* transcript was rapidly shut down and plateaued at a low and stable level (5 fold reduction after 5-7 days), RyR1 protein reduction was much slower and incomplete, being reduced by 50% only after 90 days, without further modification. This could reflect a stability of RyR1 protein to an unknown extent although *in vivo* experiments with radioactive labeling in rat muscle suggested a turnover of 8-10 days (Ferrington *et al.,* 1998, Martonosi and Halpin, 1972). Such a long stability was observed for the alpha 1 subunit of DHPR, which was knocked down with a U7-exon skipping strategy that resulted only in a 50% reduction of the protein level after 2-6 months, in presence of a remaining amount of 10-20% at the mRNA (Pietri-Rouxel *et al.,* 2010). The progressive reduction in RyR1 amount in our mouse line was associated with a progressive reduction in muscle strength, as evidenced by the direct measurement of the spontaneous ability of the animals to hang on a grid or by the evaluation of muscle tension developed during muscle electrostimulation. Our results suggest a direct correlation between the amount of RyR1 protein and the muscle strength. In our model, almost no muscle weakness was observed when RyR1 amount was above 80% of control, as already observed with the heterozygous whole body RyR1-KO mice that show a RyR1 protein amount of 85% of control and no functional alteration (Cacheux *et al.,* 2015). The drop in muscle strength in RyR1-Rec animals parallels protein amount, and when RyR1 reached 50% of control, the animals were unable to perform any of the two tests (hanging or electrostimulation). The direct study of DHPR and RyR1 function on isolated fiber demonstrated a reduction in the function of both calcium channels. The reduction in RyR1-calcium release activity was of 25%, which is similar to the reduction in RyR1 protein in the specific muscle used for the experiment (*interosseous* muscle). Surprisingly, no modification in DHPR amount or in the T-tubule density was observed, although calcium influx through DHPR was reduced by 30%. This could reflect the so-called retrograde coupling (Dirksen, 2002) by which RyR1 controls DHPR function, as observed in the RyR1-KO cells with DHPR functional alteration (Nakai *et al.,* 1996). At the structural level, all muscles were affected, and both type I (slow) and type II (fast) fibers were atrophied without any sign of basal membrane detachment (which would have reflected denervation), and with alteration in mitochondria distribution in type I fibers. A generalized disorganization of muscle structure was observed in most fibers with variable severity, from almost normal muscle structure to focal highly disorganized regions. In those highly disorganized regions, resembling the core lesions in patients, disruption of the regular alignment of the sarcomeres with abnormal mitochondria and triad localizations were observed, and the most striking feature was the presence of numerous and large stacks of membrane. Those membrane stacks, hypothetically corresponding to the close alternate apposition of SR and T-tubules, were called multiple triads (Garibaldi *et al.,* 2019), but the exact nature of each membrane type remains to be determined. Similar structures have been observed in muscle of CSQ1-KO mice (Paolini *et al.,* 2007), although smaller. The size of the SR sheets in those multiple triads is similar to the size of the SR terminal cisternae in regular triads, but the T-tubules are dilated and twice the normal size. Careful examination of those multiple triads did not allow the identification of “feets”, those electron dense structures formed by RyR1 (Block *et al.,* 1988) between SR and T-tubule membranes. So these stacks of membranes are most likely depleted of RyR1. Space size between SR and T-tubule is similar in multiple triads and in the regular triads, thus the proteins involved in the assembly of those structures are probably the same. As the RyR1 staining is uniform on transversal section, RyR1 depletion occurred most likely in all fiber types, although desmin overexpression and delocalization has been seen mainly in type I fibers. At the molecular level, some important modifications were observed, such as a 2-to 3-fold increase in CLIMP63 and in STIM1, and a massive increase in desmin (x 80). The mechanisms connecting the structural disorganizations with the overexpression of desmin, CLIMP63, STIM1 are still unclear, and require further analysis. Those modifications, both at the structural and protein levels, have been observed in mice and in human patients with RyR1 mutations resulting only in RyR1 reduction, pointing to a common pathophysiological mechanism between mouse and human. Desmin forms intermediate filaments involved in the muscle cell architecture by anchoring several structures (nucleus, mitochondria, Z-disc) to the plasma membrane at the costamers (Paulin and Li, 2004; Franck *et al.,* 2019), and alterations in desmin result in myopathies (Goldfarb and Dakalas, 2009). Desmin overexpression could reflect a direct regulation of the desmin gene by the calcium released from the sarcoplasmic reticulum. On the other hand, desmin aggregation and mislocalization leading to muscle weakness could contribute to the overall phenotype observed both in mice and in patients with RyR1 reduction. CLIMP63 is a reticulum protein, associated with the calcium release complex via its interaction with triadin (Osseni *et al.,* 2016), which is involved in the shaping of reticulum membrane and in the formation of reticulum sheets (Goyal and Blackstone, 2013). Its overexpression observed both in mice and in patients with RyR1 reduction could be associated with (and perhaps responsible for) the formation of the multiple triads observed in both cases. Along the same line, STIM1 is not only involved in the refilling of calcium stores via its interaction with ORAI1, but also in the reticulum morphogenesis via its interaction with the microtubule +TIP binding protein EB1 (Goyal and Blackstone, 2013), pointing to its possible involvement in the formation of those multiple triads. As ORAI1 undergoes the same magnitude of overexpression as STIM1, it is mostly likely that the two proteins still work together, but the functional consequences of this overexpression on the calcium entry activated by store depletion (SOCE) has to be demonstrated. The overexpression of STIM1 and ORAI1 has already been associated with SR and T-tubule remodeling during exercise (Michelucci *et al.,* 2019; Boncompagni *et al.,* 2017), resulting in the formation of stacks of few (2-4) SR membranes associated with a T-tubule, with a gap between both being smaller than the normal triadic gap. The structures observed in RyR1 deficient mouse muscle looked similar although much larger (up to 15 stacks of each membrane), and with a clear alternation of two membrane types but with the same space size between the membranes. Inhibition of autophagy has been evidenced in mice, which could explain the muscle atrophy observed, as both increase or decrease in autophagy have been described as leading to muscle atrophy (Masiero *et al.,* 2009).

To conclude, our mouse model recapitulates the main features observed in “Dusty core Disease” patients, a subgroup of Central Core Disease with only RyR1 reduction, and this model, due to a better homogeneity between the mice (same age, same sex, …) than between the patients (different severity, different age at the biopsy, different age at onset of the disease, different sex, …) and the higher number of sample available has allowed to point to modifications that could shed a new light on the pathophysiological mechanisms, only partially known, and provide new clues for therapeutic development.

## Author contributions

LP, AP, BG, DG, VJ, JF and IM designed the research studies; LP, AP, JB, BG, DG, CS, LT, MC, MB, CK, VJ, CFA, DB, DM, NR, JR, IM conducted the experiments and acquired the data; LP, AP, JB, BG, DG, VJ, CFA, JF, IM analyzed the data, IM wrote the manuscript, all the authors read and approve the manuscript

## Acknowledgments

We thank Dr Jacques Brocard for his advises with statistical analysis, Sara Virtanen for the proofreading of the manuscript, and the members of the GIN animal facility and imaging facility for their help.

## Materials and methods

### Engineering of the mouse model

The RyR1-flox mouse line has been established at the MCI/ICS (Mouse Clinical Institute, Illkirch, France; http://www-mci.u-strasbg.fr). LoxP sites were introduced by homologous recombination in ES cells on both sides of exons 9 to 11 of the *RYR1* gene. The HSA-Cre-ER^T2^ mouse line, in which the expression of the tamoxifen dependent Cre-ER^T2^ recombinase is under the control of the human skeletal muscle α-actin gene, has been described previously (Shuler *et al.,* 2005). In this transgenic line Cre-ER^T2^ is selectively expressed in skeletal muscles fibers and activated upon tamoxifen injection, and Cre-ER^T2^-mediated alteration of LoxP (floxed) target genes is skeletal muscle-specific and strictly tamoxifen dependent. The two mouse lines were on a a C57BL/6J background and intercrossed to create the RyR1^flox/flox^::HSA-Cre-ER^T2^ mouse line.

### Antibodies and reagents

Polyclonal antibody against RyR1, Trisk 95 and Trisk 51 have been previously described (Marty *et al.,*, 1994; Marty *et al.,* 2000; Osseni *et al.,* 2016), as well as antibodies against SERCA (kindly provided by Dr M.J. Moutin, Moutin *et al.,* 1994), and antibody against the FoF1-ATPase (kindly provided by Pr J. Lunardi, Lunardi *et al.,* 1989). Antibody against the alpha subunit of DHPR was from Abcam (ref ab2862), antibody against STIM1 from Merck Millipore (ref AB9870), antibody against CSQ1 was ThermoFisher (ref MA3-913), antibody against CLIMP63 from Bethyl (Rabbit anti-CKAP4 antibody, ref A302-257A), antibody against Orai1 from Alomone labs (ref ACC-062), antibody against desmin from Dako (ref M0724), polyclonal antibody against alpha-actinin from Sigma Aldrich (ref SAB4503474), antibody against p62 from Abnova (SQSTM1 monoclonal antibody clone 2C11 ref H00008878-M01), antibody against LC3A/B from Cell Signaling Technology (ref 4108), as well as antibody against mTOR and P-mTOR (ref 2972 and 2971 respectively), antibody against S6 and P-S6 (ref 2217 and 2215 respectively), rabbit mAb against GAPDH (14C10, ref 2128S). Secondary antibodies used for Western blot were labelled with HRP (Jackson Immuno Research), and the antibodies used for immunofluorescent staining were labelled with Alexa fluor 488, Cy-3 or Cy-5 (Jackson Immuno Research). Tamoxifen was from Sigma Aldrich.

### Reverse transcription-quantitative PCR (RT-qPCR)

Target gene transcript expression was measured by quantitative real-time polymerase chain reaction (RT-qPCR). Total RNA was isolated from muscle tissue using NucleoSpin RNA Set for NucleoZOL (Macherey-Nagel). RNA was quantified using a NanoDrop 1000 spectrophotometer (Thermo Scientific, Waltham, MA). RNA was reverse transcribed using iScript Reverse Transcription Supermix for RT-PCR (Bio-Rad Laboratories, Hercules, CA). Then RT-qPCR was conducted using SsoAdvanced™ Universal SYBR® Green Supermix (Bio-Rad Laboratories), and 1 μl cDNA was used to detect the transcript of interest. All reactions were run in duplicate. RT-qPCR reactions were run on an CFX96 Touch™ Real-Time PCR Detection System (Bio-Rad). The primer sequences were designed using Vector NTI software (Thermo-Fischer). The amplification steps were the following: 50°C-2min; 95°C-10min; 40 cycles composed of 95°C-15s, 60°C-30s,72°C-30s; 72°C-10min. A threefold RNA dilution series was used to determine efficiency of each qPCR assay. Efficiency of all qPCRs was between 95 and 105%. Amplification data were analyzed with CFX manager software (Bio-Rad). Expression was normalized to three endogenous controls (beta-actin, HPRT and GAPDH) using the ΔΔCt method. The primers (forward and reverse) used for PCR amplification are listed in Table 3.

**Table 3.**
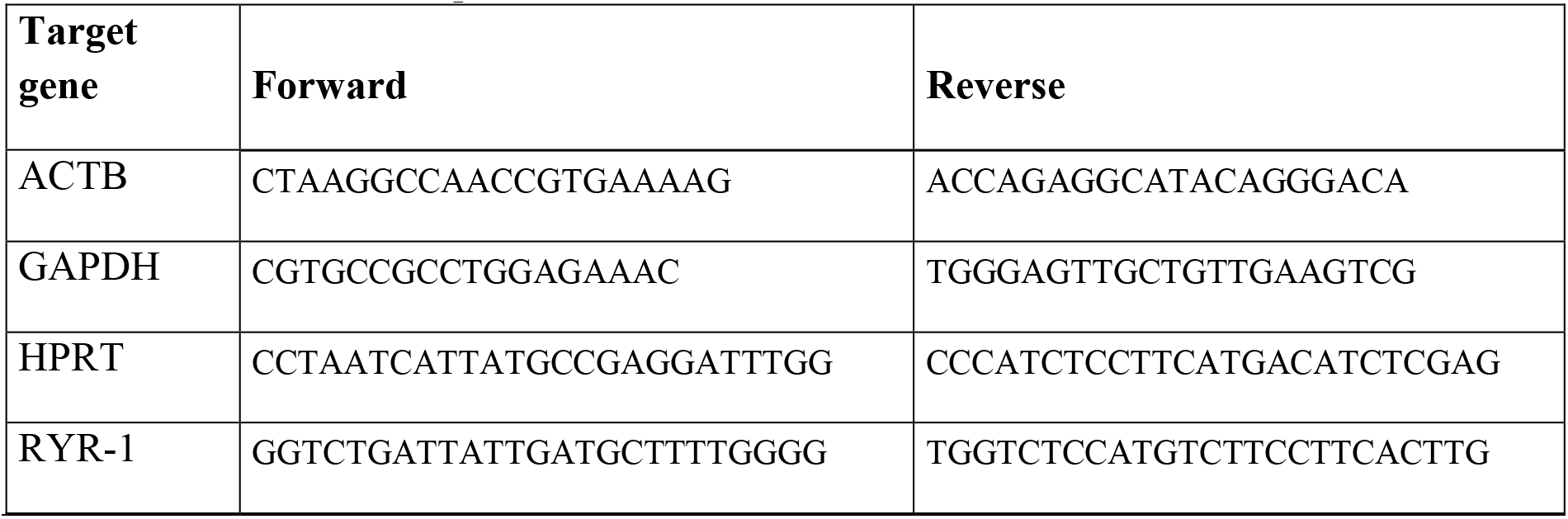
Primers for RT-qPCR

### Western blot analysis and quantification

Western blot analysis was performed on muscle homogenates as previously described (Garibaldi *et al.,* 2019). Briefly, after electrophoretic separation on a 4–20% gradient acrylamide gel (Stain-free precast gel, Biorad, France) and electrotransfer to Immobilon P (Biorad, France), the membrane was incubated with primary antibodies and then HRP-labelled secondary antibodies (Jackson ImmunoResearch Laboratories). Signal quantification was performed using a ChemiDoc Touch apparatus (Biorad, France) and the Image Lab software (Biorad). The amount of the chosen protein in each sample was corrected for differences in loading using either the amount of myosin, GAPDH or the total amount of proteins using the stain free system from Biorad, and normalized to the amount of the same protein present in the control, set to 100% as described previously (Cacheux *et al.,* 2015). Ten human controls (muscle biopsy from individuals non-affected by neuromuscular disease) of different age have been used, from 3.5 years to 64 years. For each sample (mouse and human), 2 to 3 Western blots have been performed, and the value for each sample corresponds to the mean ± SEM of the different Western blots.

### Grip test

The animals were trained 3 times on 3 different days over the week before tamoxifen injection, and they were subsequently tested once a week until muscle collection. The animals were positioned on a cross-wired surface where they could hold on with all four paws. The grid was then turned upside down, and the time during which each animal was able to stay on wires before falling was recorded, up to a maximum of 300s (5min). Three falls were allowed.

### Noninvasive investigation of gastrocnemius muscle function and bioenergetics

*Gastrocnemius* muscle anatomy and mechanical performance were investigated noninvasively and longitudinally using MR measurements in 8 controls and 6 RyR1-Rec littermates every month at D30, D60 and D90 after tamoxifen injection. Additionally, muscle energy metabolism was assessed at D60 simultaneously to mechanical performance acquisition. Anesthetized mice were placed into a home-built cradle allowing the MR investigation of the left gastrocnemius muscle function and bioenergetics in a preclinical 47/30 Biospec Avance MR scanner (Bruker, Karlsruhe, Germany) as described previously (Oddoux *et al.,* 2009; Giannesini *et al.,* 2010). Ten axial anatomic images (1-mm thickness; 0.5-mm spaced; 0.117 × 0.117 mm^2^ spatial resolution) covering the region from the knee to the ankle were acquired at rest. Mechanical performance was recorded using a dedicated ergometer during a fatiguing bout of exercise electrically induced by square-wave pulses (1-ms duration) with transcutaneous surface electrodes and consisting in 6 minutes of maximal isometric contractions repeated at a frequency of 1.7 Hz. At D60, concentrations of high-energy phosphorylated compounds and pH were continuously measured using dynamic ^31^P-MR spectroscopy during rest (6-min), exercise (6-min) and post-exercise recovery period (16-min). MR data were processed using custom-written analysis programs developed on the IDL software (Research System, Boulder, CO).

### Histological staining

Ten μm thick cryostat *Tibialis anterior* (TA) sections (Cryostar Nx70, Thermo Scientific) were stained with haematoxylin and eosin (HE), modified Gomori trichrome (GT) and reduced nicotinamide adenine dinucleotide dehydrogenase-tetrazolium reductase (NADH-TR), and observed using a Leica ICC 50 microscope.

### Electron microscopy

EDL muscles of RyR1-Rec male mice 75 days after tamoxifen injection (4.5 months old) were excised, cut into small pieces and prepared as described before (Oddoux *et al.,* 2009). The sections were observed with a Jeol JEM 1200EX II at 60 kV.

### Immunofluorescent labelling

EDL muscles were collected from CTRL and RyR1-Rec littermates 75 days after tamoxifen injection. After 15 min fixation at RT in 4% PFA diluted in PBS, the fibers were manually dissociated using needles to comb the fibers, and the triple labelling was performed as described previously (Oddoux *et al.,* 2009) using primary antibodies developed in guinea pig, rabbit or mouse, and anti-species secondary antibodies labelled with either Alexa fluor 488, Cy 3 or Cy 5. The images are representative of at least 3 animals in each group.

### Electrophysiology and confocal fluorescence imaging in isolated muscle fibers

Single fibers were isolated from the *flexor digitorum brevis* (FDB) and *interosseus* muscles following previously described procedures (Lefebvre *et al.,* 2011; Kutchuchian *et al.,* 2016, 2017). In brief, muscles were digested by incubation with collagenase followed by trituration to produce dissociated fibers. All experiments were performed at room temperature (20–22 °C). Membrane depolarizing steps of 0.5 s duration were applied from a holding potential of −80 mV to the single fibers partly insulated with silicone grease, and the resulting voltage-activated Ca^2+^ transients was measured using the dye rhod-2 introduced into the fiber by the patch pipette. The detailed electrophysiological procedures are described in Supplementary methods.

### Ethics Approval

All procedures using animals were approved by the institutional ethics committee (CEEA-GIN 04, N_134) and followed the guidelines from Directive 2010/63/EU of the European Parliament on the protection of animals used for scientific purposes. This study adheres to the ARRIVE guidelines. Experimental procedures for human studies were approved by the institutional ethics committee and performed in accordance with ethical standards laid down in the Declaration of Helsinki. All biopsies were collected following informed consent.

### Statistics

The statistical analysis has been done with GraphPad Prism 6.0 software. The number of samples and the name of the parametric test applied are indicated in each figure legend. Results are considered as significant when *p < 0.05*, the exact value for *p* being indicated in the text or figure legends, and significant results are labeled * on the graphs whatever the exact *p* value. All data are shown as mean ± SEM.

### Data availability

The data that support the findings of this study are available from the corresponding author, upon reasonable request.

