## supplementary material (methods, figures S1 to S6) for "*In vivo* RyR1 reduction in muscle triggers a core-like myopathy"

### Supplementary method:

*Electrophysiology and confocal fluorescence imaging in isolated muscle fibers*
We used previously described procedures (Lefebvre *et al*., 2011; Kutchukian *et al*., 2016, 2017). All solutions were adjusted to pH 7.20. All experiments were performed at room temperature (20-22° C). F*lexor digitorum brevis* and interosseus muscles were isolated and incubated with collagenase (Sigma, type 1) for 60 min at 37°C in the presence of Tyrode solution containing (in mM): 140 NaCl, 5 KCl, 2.5 $CaCl_2$, 2 $MgCl_2$, 10 HEPES. Single fibers were obtained by mechanical trituration of a given muscle within a 50 mm wide culture µ-dish (Ibidi, Planegg / Martinsried, Germany).
For all experiments requiring voltage-clamp, the bottom of the µ-dish was first coated with a thin layer of silicone and then filled with culture medium containing 10% fetal bovine serum (MI199; Eurobio, France); fibers from the interosseus muscles were isolated with the trituration procedure and were embedded within silicone so that only a portion of the fiber extremity remained out, in contact with the extracellular medium. The culture medium was then replaced by the standard extracellular solution containing (in mM) 140 TEA-methane-sulfonate, 2.5 $CaCl_2$, 2 $MgCl_2$, 1 4-aminopyridine, 10 HEPES and 0.002 tetrodotoxin, and, for measurements of fluo-4 FF $Ca^{2+}$ transients, 0.05 N-benzyl-p-toluene sulfonamide (BTS) to block contraction. Voltage-clamp was achieved with a micropipette filled with an intracellular-like medium containing (in mM) 120 K-glutamate, 5 $Na_2$-ATP, 5 $Na_2$-phosphocreatine, 5.5 $MgCl_2$, 5 glucose, 5 HEPES. For measurements of fluo-4FF $Ca^{2+}$ transients it also contained 0.1 fluo-4 FF. For measurements of rhod-2 $Ca^{2+}$ transients it also contained 15 EGTA, 6 $CaCl_2$ and 0.1 rhod-2. The tip of the pipette was inserted into the silicone-insulated part of the fiber and was gently crushed against the bottom of the chamber to ease intracellular dialysis and reduce series resistance. The fiber interior was left to

equilibrate for 30 min before starting measurements. The pipette was connected to an RK-400 patch-clamp amplifier (Bio-Logic, Claix, France) used in whole-cell voltage-clamp configuration, in combination with an analog-digital converter (Digidata 1440A, Axon Instruments, Foster City, CA) controlled by pClamp 9 software (Axon Instruments). Analog compensation was adjusted to further decrease the effective series resistance. The holding voltage was set to -80 mV.

DHPR $Ca^{2+}$ current was measured in response to 0.5 s-long depolarizing steps of increasing amplitude. The passive linear component of the current was removed by subtracting the adequately scaled value of the steady current measured in response to a -20 mV step. Values for peak $Ca^{2+}$ current were further corrected by the linear slope of the I/V relationship between -50 and -30 mV. Peak $Ca^{2+}$ current values were normalized to the fiber capacitance (current density). The voltage dependence of the peak $Ca^{2+}$ current density was fitted with the following equation:

$I(V) = G_{max}(V-V_{rev})/(1+\exp(V_{0.5}-V)/k])$

with $I(V)$ the peak current density at the command voltage $V$, $G_{max}$ the maximum conductance, $V_{rev}$ the apparent reversal potential, $V_{0.5}$ the half-activation potential and $k$ the steepness factor.

Confocal imaging was conducted with a Zeiss LSM 5 Exciter microscope equipped with a 63× oil immersion objective (numerical aperture 1.4). For detection of $Ca^{2+}$ transients with rhod-2 and fluo-4 FF fluorescence, excitation was from the 543 nm line of a HeNe laser and from the 488 nm line of an Argon laser, respectively, and fluorescence was collected above 560 nm and above 505 nm, respectively. Imaging was performed with the line-scan mode ($x,t$) of the system and quantified as $F/F_0$ where $F_0$ is the baseline fluorescence. The rate of SR $Ca^{2+}$ release ($d[Ca_{Tot}]/dt$) was calculated from the rhod-2 $Ca^{2+}$ transients as previously described (Lefebvre *et al.*, 2011; Kutchukian *et al.*, 2016). In each fiber, the voltage-dependence of the peak rate of $Ca^{2+}$ release was fitted with a Boltzmann function:

$d[Ca_{Tot}]/dt\ (V)= Max\ d[Ca_{Tot}]/dt\ /(1+\exp[(V_{0.5}-V)/k])$

$Max\ d[Ca_{Tot}]/dt$ corresponding to the maximum rate of $Ca^{2+}$ release, $V_{0.5}$ to the mid-activation voltage and $k$ to the steepness factor.

For imaging the T-tubule network with di-8-anepps, fluorescence was collected above 505 nm with 488 nm excitation. For this, isolated *flexor digitorum brevis* fibers were incubated for 30 minutes in the presence of 10 µM di-8-anepps in Tyrode solution. The T-tubule density was estimated as described previously (Kutchukian *et al.*, 2017). In brief, 2 *x,y* images were taken at distinct locations along each fiber. Analysis was carried out with the ImageJ software (National Institute of Health): automatic thresholding with the Otsu method was used to create a binary image of the area occupied by t-tubules. The *skeletonize* function was then used to delineate the t-tubule network. T-tubule density was expressed as the percent of positive pixels within the region (referred to as T-tubule density index).

### Supplementary Figures

**Figure S1: Evaluation of RyR1 transcript, protein amounts and muscle weight at D75 in different muscles**

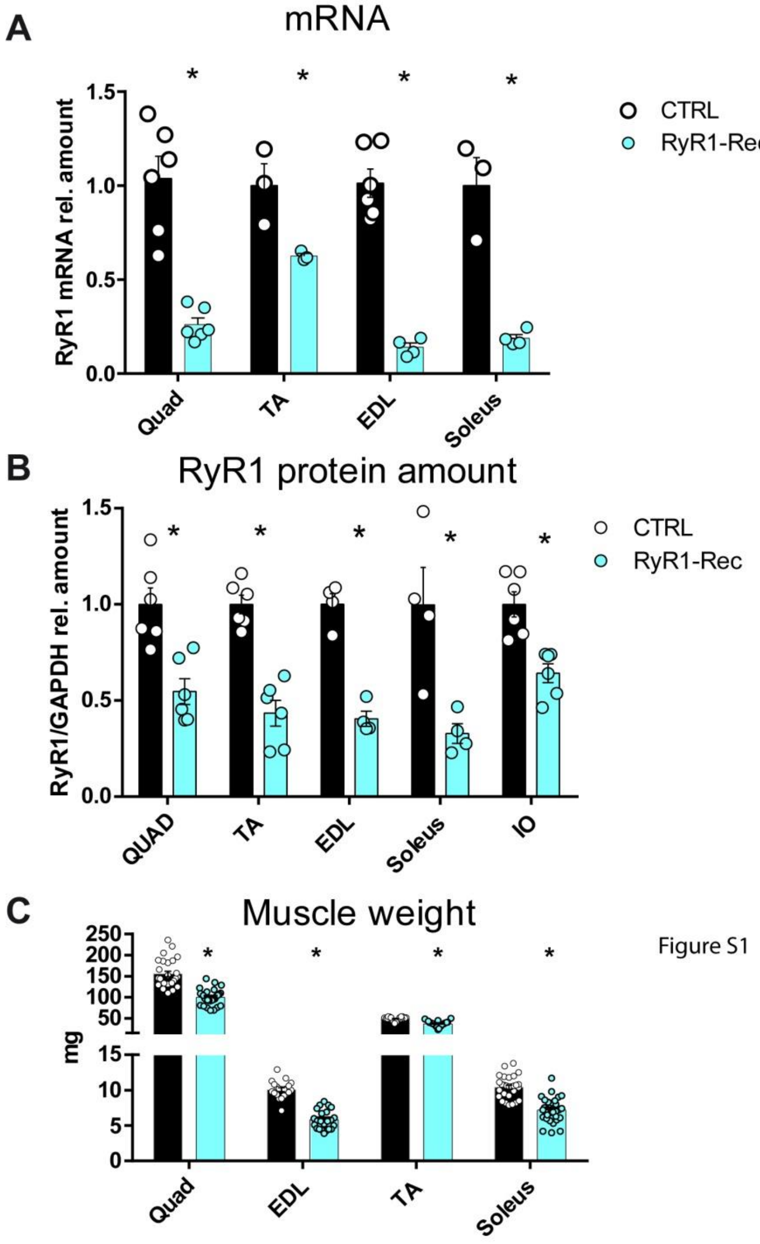


Black bars represent the CRTL animals and blue bars the RyR1-Rec animals. Data are presented as mean ± SEM of *n=3-6* animals. Statistical analysis: t-test with Holm-Sidak method for multiple comparisons between CTRL and RyR1-Rec for the same muscle.* $p<0.05$.

(A) The relative amount of RyR1-mRNA compared to beta-actin, HPRT and GAPDH as reference genes was evaluated using RT-q-PCR in different muscles from WT and RyR1-Rec animals. The relative amount was normalized to CTRL at each time. The quantification was performed using the ΔΔCt method on *n=3-6* animals in each group. (B) The relative amount of RyR1 compared to GAPDH was evaluated using quantitative Western blot in different muscle homogenates of *n=3-6* different mice. The mean of protein amount in CTRL littermate was set to 1. (C) The weight of the different muscles was measured at D75 on 20-30 muscles in each group.

**Figure S2: *In vivo* bioenergetics analysis of gastrocnemius function in D60 mice**

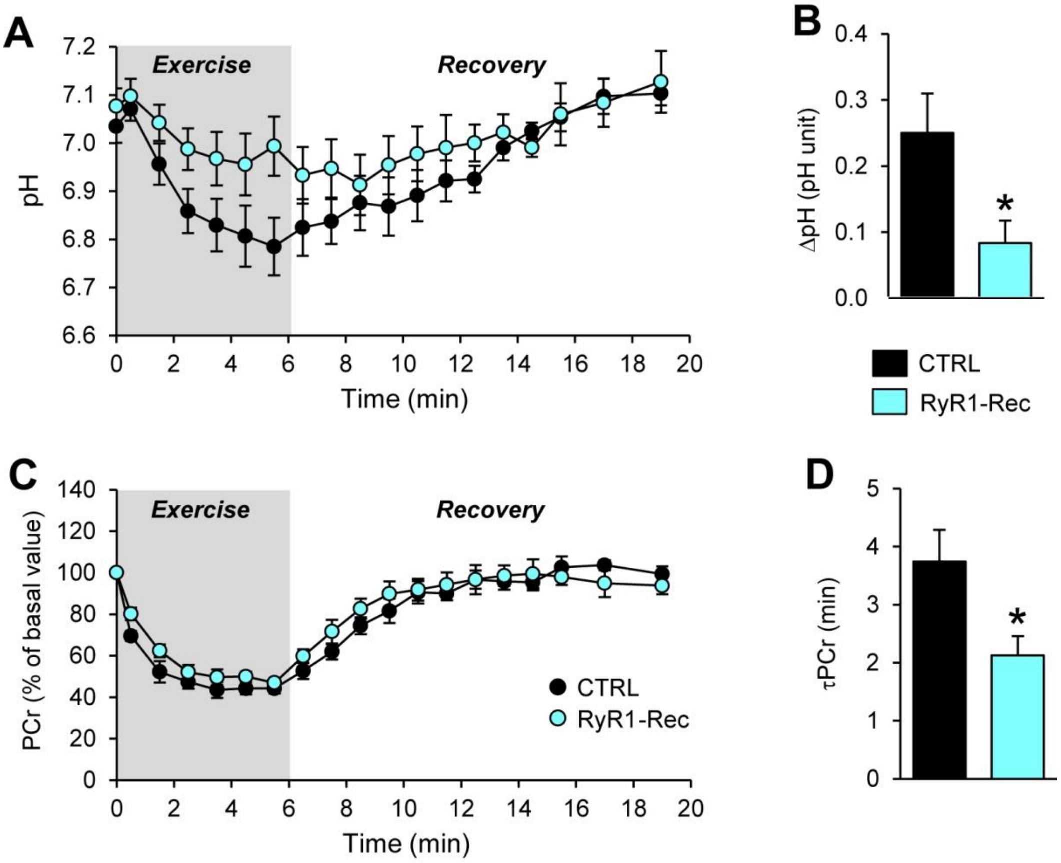


Changes in pH (A) and [PCr] (C) were measured *in vivo* at D60 in exercising (simultaneously to mechanical performance acquisition) and recovering muscle of 8 CTRL and 6 RyR1-Rec mice. For panels A and C, the first time-point (t = 0) indicates the basal value. In order to determine the time constant of post-exercise phosphocreatine resynthesis (τPCr, an *in vivo* index of mitochondrial capacity), the time course of phosphocreatine (PCr) during the post-exercise period was fitted to a monoexponential function with a least mean-squared algorithm: $\tau PCr = -t/\ln(PCr_t/\Delta PCr)$, where ΔPCr is the extent of PCr depletion measured at the start of the recovery period. For RyR1-Rec mice, the extent of acidosis at the exercise end was lower (B), whereas the PCr recovery time constant (τPCr) was shorter (D) when compared to CTRL. Data are means ± SEM. * $p < 0.05$ *vs*. CTRL animals (unpaired two-tailed Student's t-tests).

**Figure S3: Cytosolic $Ca^{2+}$ removal capabilities of isolated muscle fibers.**

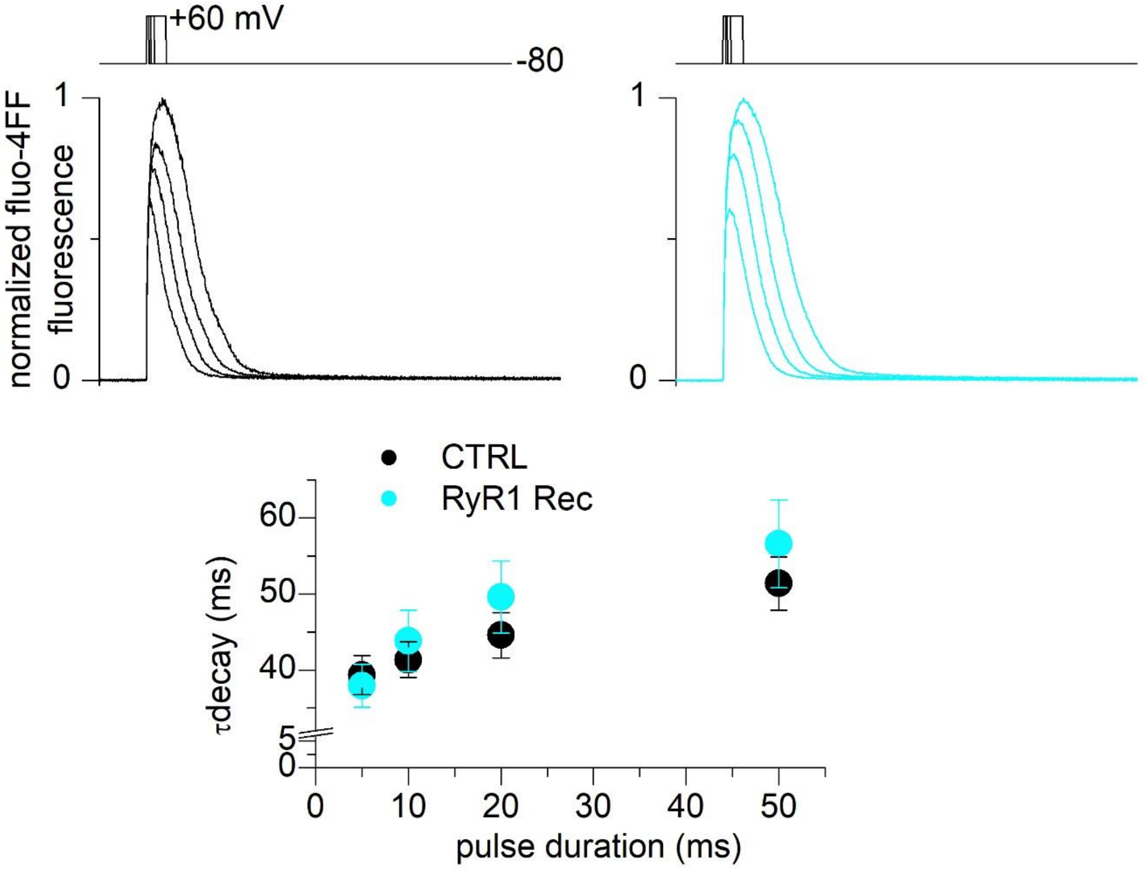


Fluo-4FF $Ca^{2+}$ transients elicited by voltage-clamp depolarizing pulses of 5, 10, 20 and 50 ms duration from -80 to +60 mV in a CTRL muscle fiber (left) and in a RyR1-Rec muscle fiber (right). Transients were normalized to the peak amplitude of the transient in response to the 50 ms-long pulse. In each fiber, the decaying phase of the transients, after the end of the pulses, was fitted with a single exponential function. The graph shows mean values for the time constant of decay in the two groups of fibers. Data are from 11 fibers from 5 CTRL mice and from 14 fibers from 5 RyR1-Rec mice, respectively.

**Figure S4: Histological staining with Gomori trichrome of TA sections of D75 mice**

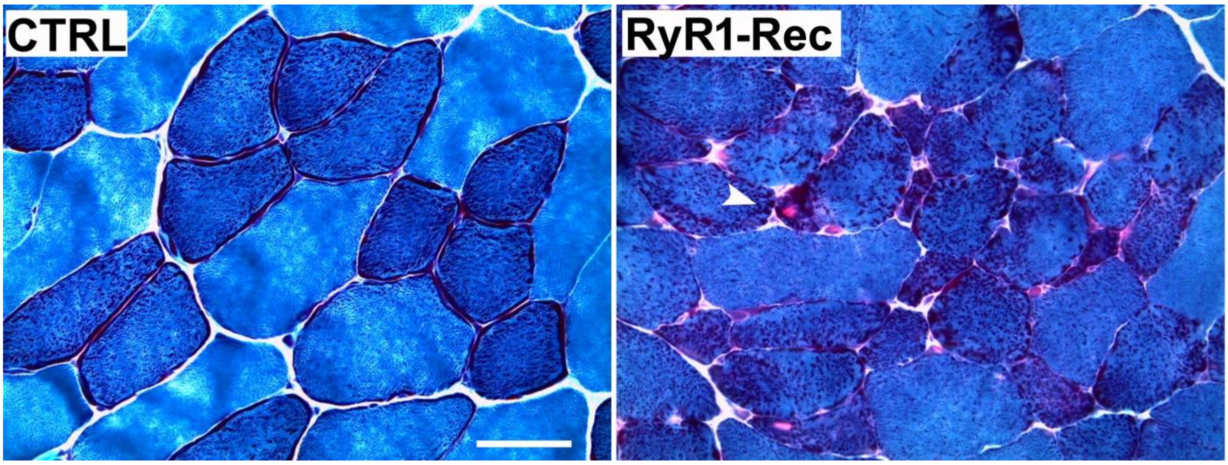


TA sections from CTRL and RyR1-Rec D75 mice were stained with Gomori trichrome. Accumulation of reddish stained material corresponding to mitochondria can be seen in some fibers of RyR1-Rec muscle (arrow). Bar 50µm

**Figure S5: Morphological measurement of triads' features**

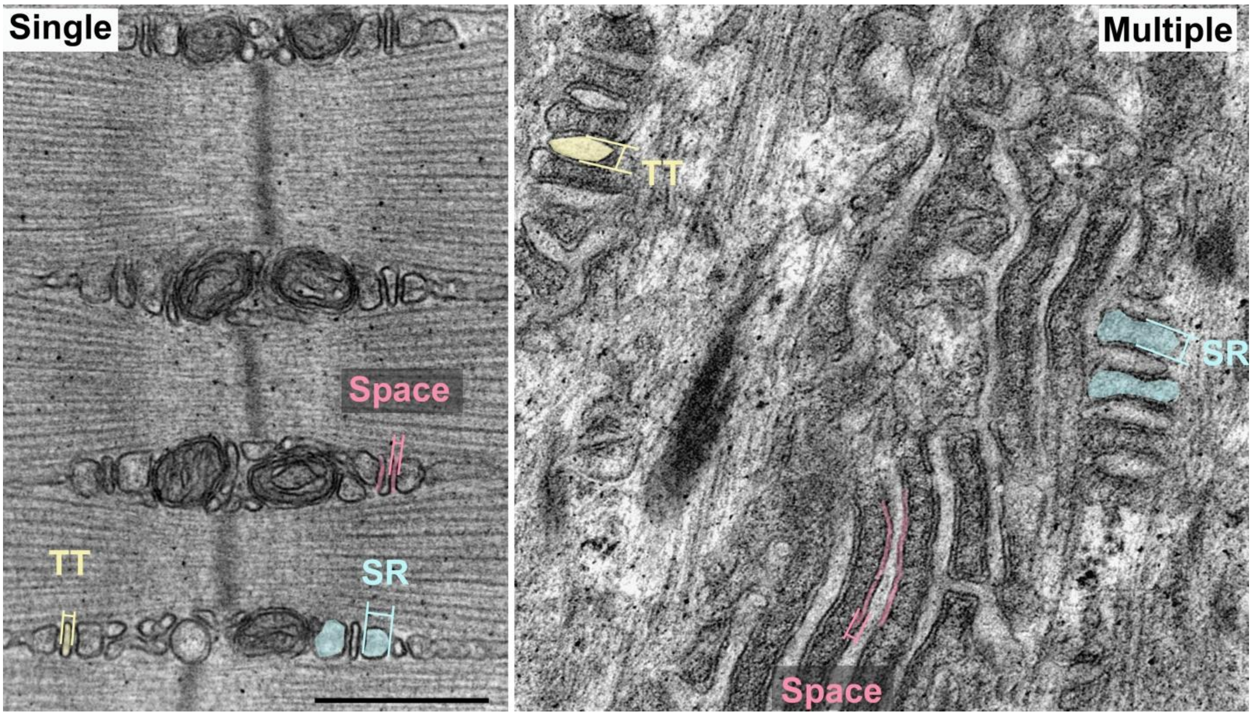


Single triads from CRTL animals or multiple triads from RyR1-Rec animals were analyzed, and the thickness of T-tubules (one TT is colored in yellow), sarcoplasmic reticulum sheets (SR, two are colored in blue) and the size of the space between T-tubule and SR membrane were measured as presented on the two representative images. The two images are at the same scale, bar 500nm.

**Figure S6: Immunofluorescent staining of muscle fibers.**

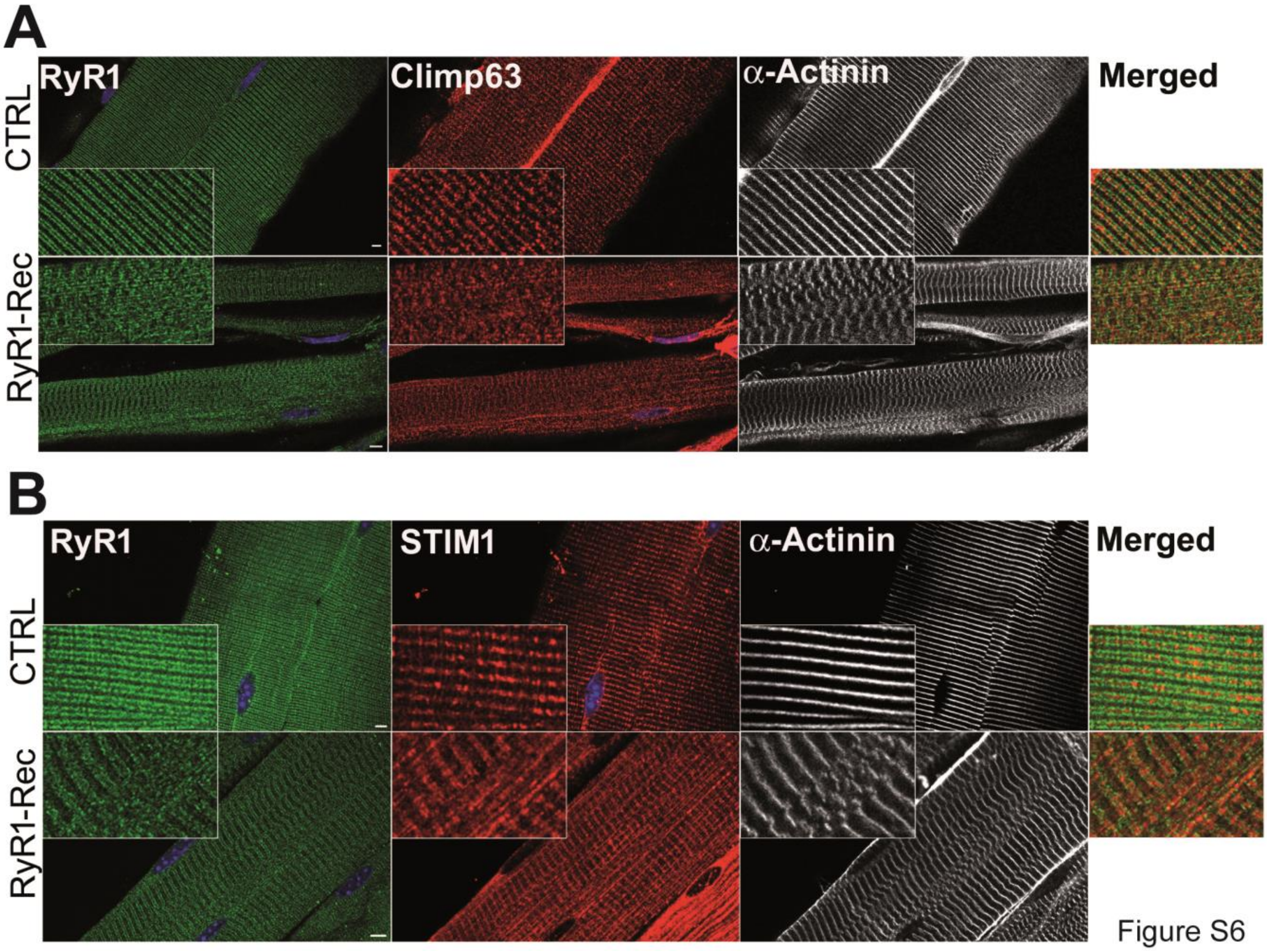


(A) Dissociated EDL fibers from D75 CTRL and RyR1-Rec mice were stained with antibodies against RyR1 (green), CLIMP63 (red) and alpha-actinin (white). Bar 2µm. (B) Dissociated EDL fibers from D75 CTRL and RyR1-Rec mice were stained with antibodies against RyR1 (green), STIM1 (red) and alpha-actinin (white). Bar 2µm
